# Imbalance of peptidoglycan biosynthesis alters the cell surface charge of *Listeria monocytogenes*

**DOI:** 10.1101/2022.09.13.507711

**Authors:** Lisa Maria Schulz, Patricia Rothe, Sven Halbedel, Angelika Gründling, Jeanine Rismondo

## Abstract

The bacterial cell wall is composed of a thick layer of peptidoglycan and cell wall polymers, which are either embedded in the membrane or linked to the peptidoglycan backbone and referred to as lipoteichoic acid (LTA) and wall teichoic acid (WTA), respectively. Modifications of the peptidoglycan or WTA backbone can alter the susceptibility of the bacterial cell towards cationic antimicrobials and lysozyme. The human pathogen *Listeria monocytogenes* is intrinsically resistant towards lysozyme, mainly due to deacetylation and *O*-acetylation of the peptidoglycan backbone via PgdA and OatA. Recent studies identified additional factors, which contribute to the lysozyme resistance of this pathogen. One of these is the predicted ABC transporter, EslABC. An *eslB* mutant is hyper-sensitive towards lysozyme, likely due to the production of thinner and less *O*-acetylated peptidoglycan. Using a suppressor screen, we show here that suppression of *eslB* phenotypes could be achieved by enhancing peptidoglycan biosynthesis, reducing peptidoglycan hydrolysis or alterations in WTA biosynthesis and modification. The lack of EslB also leads to a higher negative surface charge, which likely stimulates the activity of peptidoglycan hydrolases and lysozyme. Based on our results, we hypothesize that the portion of cell surface exposed WTA is increased in the *eslB* mutant due to the thinner peptidoglycan layer and that latter one could be caused by an impairment in UDP-*N*-acetylglucosamine (UDP-Glc*N*Ac) production or distribution.

## 1. INTRODUCTION

In recent years, antibiotic resistance became a serious threat for public health. One of the main targets of currently available antibiotics is the bacterial cell wall (Fig. 1). In Gram-positive bacteria, the cell wall consists of a thick layer of peptidoglycan and cell wall polymers, which are either tethered to the cell membrane or covalently linked to peptidoglycan and referred to as lipoteichoic (LTA) and wall teichoic acid (WTA), respectively. Peptidoglycan biosynthesis starts in the cytoplasm by the conversion of UDP-*N*-acetylglucosamine (UDP-Glc*N*Ac) to UDP-*N*-acetylmuramic acid (UDP-Mur*N*Ac) via MurA and MurB. The activity of the UDP-Glc*N*Ac 1-carboxyvinyltransferase MurA can be specifically inhibited by the antibiotic fosfomycin (Fig. 1)(Kahan et al., 1974). In the subsequent steps, which are depicted in Figure 1, the peptidoglycan precursor lipid II is synthesized in the cytoplasm and transported across the membrane by the flippase MurJ (Meeske et al., 2015; Ruiz, 2008; Sham et al., 2014). Lipid II is then incorporated into the growing glycan strand by members of the SEDS (shape, elongation, division, sporulation) protein family, namely RodA and FtsW, which act in concert with class B penicillin binding proteins (PBPs) that have a transpeptidase activity (Cho et al., 2016; Leclercq et al., 2017; Taguchi et al., 2019). Recent findings led to the conclusion that class A PBPs, which possess a glycosyltransferase and a transpeptidase domain, are mainly involved in filling gaps and/or repair defects in the peptidoglycan mesh, rather than being the main enzymes involved in peptidoglycan polymerization and crosslinking (Cho et al., 2016; Dion et al., 2019; Vigouroux et al., 2020). The ability of bacterial cells to elongate and divide depends on the activity of peptidoglycan hydrolases. Two DL-endopeptidase, CwlO and LytE, are essential for cell elongation in *Bacillus subtilis*. In addition to its role in cell elongation, LytE is also involved in cell separation (Carballido-López et al., 2006; Ohnishi et al., 1999; Vollmer et al., 2008). Both enzymes cleave the peptide bond between d-glutamic acid and *meso*-diamino pimelic acid of the peptidoglycan peptide, thereby allowing the insertion of new peptidoglycan material (Yamaguchi et al., 2004). Lack of CwlO results in cell shortening, while absence of CwlO and LytE is lethal (Domínguez-Cuevas et al., 2013; Hashimoto et al., 2012). Peptidoglycan biosynthesis and hydrolysis need to be tightly regulated to prevent cell lysis. In *B. subtilis*, this regulation is partly achieved by controlling expression of *cwlO* and *lytE* via the essential two-component system WalRK, whose expression is for instance induced during heat stress (Bisicchia et al., 2007; Dubrac et al., 2008; Takada et al., 2018). On the other hand, activity of CwlO depends on a direct protein-protein interaction with the ABC transporter FtsEX (Meisner et al., 2013). The FtsEX-dependent regulation of *B. subtilis* CwlO further depends on the presence of two cofactors, SweC and SweD (Brunet et al., 2019; Rismondo and Schulz, 2021).

**Figure 1:**
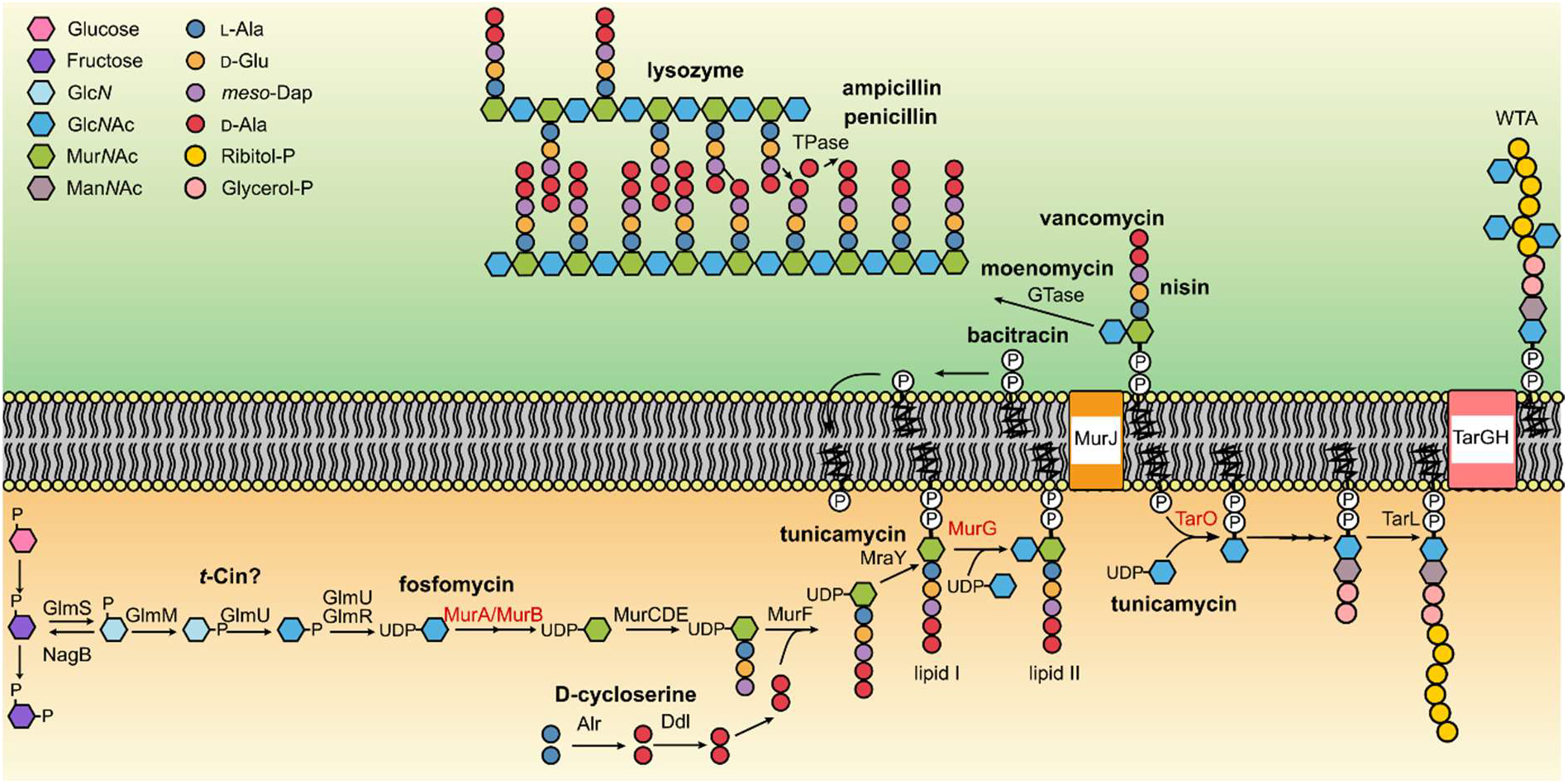
Schematic of the UDP-Glc*N*Ac, peptidoglycan and wall teichoic acid biosynthesis pathway. UDP-Glc*N*Ac, which is required for peptidoglycan and wall teichoic acid (WTA) biosynthesis, is synthesized from Fructose-6-phosphate in a four-step reaction catalyzed by GlmS, GlmM, GlmU and GlmR. MurA, an UDP-Glc*N*Ac-1-carboxyvinyltransferase catalyzing the first step of peptidoglycan biosynthesis, and MurB convert UDP-Glc*N*Ac to UDP-Mur*N*Ac. Lipid II is produced in subsequent steps, which are performed by MurCDEF, Alr, Ddl, MraY and MurG, flipped across the membrane by MurJ and inserted into the growing glycan strand by glycosyltransferases (GTases). Finally, peptidoglycan is crosslinked by the action of transpeptidases (TPases) (Pazos and Peters, 2019). UDP-Glc*N*Ac also serves as a substrate of TarO, the first enzyme of the WTA biosynthesis pathway. After the WTA polymer is synthesized by a subset of enzymes, it is transported across the membrane via TarGH (Brown et al., 2013) and, in case of *L. monocytogenes* 10403S, decorated with Glc*N*Ac residues (Shen et al., 2017). UDP-Glc*N*Ac-consuming enzymes are labelled in red. Antibiotics targeting different steps of the UDP-Glc*N*Ac, peptidoglycan and WTA biosynthesis pathways or degrade peptidoglycan are depicted in bold (Campbell et al., 2011; Pensinger et al., 2021; Sarkar et al., 2017).

UDP-Glc*N*Ac is synthesized from fructose-6-phosphate via a four-step reaction catalyzed by GlmS, GlmM, GlmU and GlmR and serves as a substrate of MurA. Recent findings suggest that the UDP-Glc*N*Ac biosynthetic pathway can be inhibited by *trans*-cinnamaldehyde (*t*-Cin) (Fig. 1)(Pensinger et al., 2021; Sun et al., 2021). UDP-Glc*N*Ac is also consumed by TarO/TagO, which catalyzes the first committed step for the synthesis of WTA (Soldo et al., 2002). The activity of TarO/TagO can be blocked by tunicamycin, which also affects the activity of MraY at high concentrations (Campbell et al., 2011; Hakulinen et al., 2017; Price and Tsvetanova, 2007; Watkinson et al., 1971). In *L. monocytogenes* 10403S, WTA is composed of a glucose-glucose(Glc-Glc)-glycerol phosphate-Glc*N*Ac-*N*-acetylmannosamine (Man*N*Ac) linker unit and an anionic ribitol phosphate backbone, which can be modified with rhamnose, Glc*N*Ac and d-alanine residues (Shen et al., 2017). The modification of WTA with positively charged d-alanine residues helps to mask the negative charge of the ribitol phosphate backbone, thereby conferring resistance towards cationic antimicrobial peptides and lysozyme (Brown et al., 2013; Vadyvaloo et al., 2004).

Lysozyme is an enzyme that cleaves the β-1,4-glycosidic bond between Mur*N*Ac and Glc*N*Ac of the bacterial peptidoglycan backbone and is found in human body fluids such as tears, saliva and mucus. *L. monocytogenes* is intrinsically resistant towards lysozyme, which is mainly achieved by modifications of the peptidoglycan. PgdA, an *N*-deacetylase, and OatA, an *O*-acetyltransferase, deacetylate and acetylate the Glc*N*Ac and Mur*N*Ac residues, respectively (Aubry et al., 2011; Boneca et al., 2007). In addition to peptidoglycan modifying enzymes, lysozyme resistance of *L. monocytogenes* is affected by the activity of the predicted carboxypeptidase PbpX, the non-coding RNA Rli31 and the transcription factor DegU (Burke et al., 2014). Interestingly, it was also been observed that the lack of components of the putative ABC transporter EslABC leads to a strong reduction in lysozyme resistance (Burke et al., 2014; Durack et al., 2015; Rismondo et al., 2021b). Recently, we could show that the absence of EslB, one of the transmembrane proteins of the EslABC transporter, resulted in the production of a thinner peptidoglycan layer and a reduction in O-acetylation of the peptidoglycan, which likely contributes to the reduced lysozyme resistance. Additionally, we observed that the *eslB* mutant is unable to grow in media containing high sugar concentrations and that the strain has a cell division defect (Rismondo et al., 2021b; Rismondo and Schulz, 2021).

In the current study, we used a suppressor screen to gain further insights into the role of EslABC on the physiology of *L. monocytogenes*. This screen revealed that phenotypes of the *eslB* mutant can either be suppressed by enhancing peptidoglycan biosynthesis, reducing peptidoglycan hydrolysis or by altering WTA production or modification. Using a cytochrome C assay, we further demonstrate that the lack of EslB manifests in a higher negative surface charge, which likely affects the activity of peptidoglycan hydrolases and provides an additional explanation for the increased lysozyme sensitivity of the *eslB* mutant.

## 2. MATERIALS AND METHODS

### 2.1 Bacterial Strains and growth conditions

All strains and plasmids used in this study are listed in Table S1. *Escherichia coli* strains were grown in Luria-Bertani (LB) medium and *Listeria monocytogenes* strains in brain heart infusion (BHI) medium at 37°C unless otherwise stated. Where required, antibiotics and supplements were added to the medium at the following concentrations: for *E. coli* cultures, ampicillin (Amp) at 100 μg ml^−1^, kanamycin (Kan) at 30 μg ml^−1^, and for *L. monocytogenes* cultures, chloramphenicol (Cam) at 10 μg ml^−1^, erythromycin (Erm) at 5 μg ml^−1^, Kan at 30 μg ml^−1^, nalidixic acid (Nal) at 30 μg ml^−1^, streptomycin (Strep) at 200 μg ml^−1^ and IPTG at 1 mM.

### 2.2 Strain and plasmid construction

All primers used in this study are listed in Table S2. For the construction of pIMK3-*murA*, pIMK3-*glmR*, pIMK3-*glmU* and pIMK3-*glmM*, the *murA*, *glmR*, *glmU* and *glmM* genes were amplified using primer pairs JR90/JR91, JR134/JR135, JR136/JR137 and JR138/JR139, respectively, cut with NcoI and SalI and ligated into plasmid pIMK3 that had been cut with the same enzymes. The resulting plasmids pIMK3-*murA*, pIMK3-*glmR*, pIMK3-*glmU* and pIMK3-*glmM* were recovered in *E. coli* XL1-Blue yielding strains EJR52, EJR106, EJR107 and EJR108. Next, plasmids pIMK3-*murA*, pIMK3-*glmR*, pIMK3-*glmU* and pIMK3-*glmM* were transformed into *E. coli* S17-1 yielding strains EJR59, EJR132, EJR133 and EJR134. Strain EJR59 was used as a donor strain to transfer plasmid pIMK3-*murA* by conjugation into *L. monocytogenes* strains 10403S (ANG1263) and 10403SΔ*eslB*_(*2*)_ (ANG5662) using a previously described method (Lauer et al., 2002). This resulted in the construction of strains 10403S pIMK3-*murA* (LJR26) and 10403SΔ*eslB*_(*2*)_ pIMK3-*murA* (LJR27), in which the expression of *murA* is under the control of an IPTG-inducible promoter. Strains carrying the empty pIMK3 vector were used as controls. For this purpose, pIMK3 was transformed into *E. coli* S17-1 yielding strain EJR58. S17-1 pIMK3 was subsequently used as a donor strain to transfer plasmid pIMK3 by conjugation into *L. monocytogenes* strains 10403S (ANG1263) and 10403SΔ*eslB*_(*2*)_ (ANG5662), which resulted in the construction of strains 10403S pIMK3 (LJR24) and 10403SΔ*eslB*_(*2*)_ pIMK3 (LJR25). For the construction of pIMK3-*glmS*, the *glmS* gene was amplified using primer pair JR140/JR141. The resulting PCR product was cut with *BamH*I and *Xma*I and ligated into plasmid pIMK3 that had been cut with the same enzymes. Plasmid pIMK3-*glmS* was recovered in *E. coli* XL1-Blue and subsequently transformed into *E. coli* S17-1 yielding strains EJR112 and EJR135, respectively. Strains EJR132, EJR133, EJR134 and EJR135 were used to transfer plasmids pIMK3-*glmR*, pIMK3-*glmU*, pIMK3-*glmM* and pIMK3-*glmS* by conjugation into *L. monocytogenes* strain 10403SΔ*eslB*_(*2*)_ (ANG5662) yielding strains 10403SΔ*eslB*_(*2*)_ pIMK3-*glmR* (LJR63), 10403SΔ*eslB*_(*2*)_ pIMK3-*glmU* (LJR64), 10403SΔ*eslB*_(*2*)_ pIMK3-*glmM* (LJR65) and 10403SΔ*eslB*_(*2*)_ pIMK3-*glmS* (LJR66).

For the construction of a markerless deletion of *cwlO* (*lmrg_01743*), 1-kb DNA fragments up- and downstream of *cwlO* were amplified by PCR with primers LMS106/107 and LMS104/105. The resulting PCR products were fused in a second PCR using primers LMS105/106, the product cut with *BamH*I and *Kpn*I and ligated into pKSV7 that had been cut with the same enzymes. The resulting plasmid pKSV7-Δ*cwlO* was recovered in *E. coli* XL1-Blue yielding strain EJR63. Plasmid pKSV7-Δ*cwlO* was subsequently transformed into *L. monocytogenes* 10403S and *cwlO* deleted by allelic exchange according to a previously published method (Camilli et al., 1993) yielding strain 10403SΔ*cwlO* (LJR37). Since attempts of producing electrocompetent *cwlO* cells were unsuccessful, plasmid pIMK3-*cwlO* was first conjugated into strain LJR37, which allows for IPTG-inducible expression of *cwlO*, and resulting in the construction of strain LJR103. For the construction of pIMK3-*cwlO*, *cwlO* was amplified using the primer pair LMS226 and LMS227. The fragment was cut with *Nco*I and *BamH*I, ligated into pIMK3 that had been cut with the same enzymes and recovered in *E. coli* XL1-blue and S17-1, yielding strains EJR114 and EJR115, respectively. To generate a *cwlO eslB* double mutant, plasmid pKSV7-Δ*eslB* was first transformed into *L. monocytogenes* strain LJR103 resulting in strain LJR114. The *eslB* gene was then deleted by allelic exchange. This resulted in the construction of strain 10403SΔ*cwlO* Δ*eslB* pIMK3-*cwlO* (LJR119).

For the construction of promoter-*lacZ* fusions, the plasmid pAC7 was used. The promoter region of the *dlt* operon was amplified from genomic DNA of the *L. monocytogenes* wildtype 10403S and the *eslB* suppressor strain ANG5746 containing a mutation 31 bp upstream of the ATG start codon, using the primer pair LMS93 and LMS94. The PCR products were digested with BamHI and EcoRI and ligated into pAC7. The resulting plasmids pAC7-P_*dlt*_ and pAC7-P_*dlt*_* were recovered in XL1 Blue resulting in strains EJR50 and EJR51. Subsequently, both plasmids were integrated into the *amyE* site of the *B. subtilis* wildtype strain 168, resulting in strains BLMS4 and BLMS5, respectively.

### 2.3 Generation of *eslB* suppressors and whole genome sequencing

For the generation of *eslB* suppressors, overnight cultures of three independently generated *L. monocytogenes eslB* mutants, strains 10403SΔ*eslB*_(*1*)_ (ANG4275), 10403SΔ*eslB*_(*2*)_ (ANG5662) and 10403SΔ*eslB*_(*3*)_ (ANG5685), were adjusted to an OD_600_ of 1. 100 μl of 10^−1^ and 10^−2^ dilutions of these cultures were plated on BHI plates containing (A) 0.5 M sucrose and 0.025 μg ml^−1^ penicillin, (B) 0.5 M sucrose and 0.05 μg ml^−1^ penicillin or (C) 100 μg ml^−1^ lysozyme, conditions under which the *eslB* mutant is unable to grow while the *L. monocytogenes* wildtype strain 10403S can grow. The plates were incubated at 37°C overnight and single colonies were re-streaked on BHI plates. This procedure was repeated at least three independent times per condition. The genome sequence of a selection of *eslB* suppressors was determined by whole genome sequencing (WGS) using an Illumina MiSeq machine and a 150 paired end Illumina kit as described previously (Rismondo et al., 2021b). The reads were trimmed, mapped to the *L. monocytogenes* 10403S reference genome (NC_017544) and single nucleotide polymorphisms (SNPs) with a frequency of at least 90% identified using CLC workbench genomics (Qiagen) and Geneious Prime^®^ v.2021.0.1. The whole genome sequencing data were deposited at the European Nucleotide Archive (ENA) under accession number PRJEB55822.

### 2.4 Determination of resistance towards antimicrobials

For the disk diffusion assays, overnight cultures of the indicated *L. monocytogenes* strains were adjusted to an OD_600_ of 0.1. 100 μl of cultures were spread on BHI agar plates using a cotton swap. Plates contained 1 mM IPTG were indicated. 6 mm filter disks were placed on top of the agar surface, soaked with 20 μl of the appropriate antibiotic stock solution and the plates were incubated at 37°C. The diameter of the inhibition zone was measured the next day. The following stock solutions were used: 50 mg ml^−1^ fosfomycin, 1 mg ml^−1^ *t*-Cin, 5 mg ml^−1^ tunicamycin, 15 mg ml^−1^ d-cycloserine, 60 mg ml^−1^ nisin, 30 mg ml^−1^ vancomycin, 5 mg ml^−1^ moenomycin, 1 mg ml^−1^ penicillin, 10 mg ml^−1^ ampicillin and 250 mg ml^−1^ bacitracin.

### 2.5 Spot plating assays

Overnight cultures of *L. monocytogenes* strains were adjusted to an OD_600_ of 1 and serially diluted to 10^−6^. 5 μl of each dilution were spotted on BHI agar plates, BHI agar plates containing 0.5 M sucrose and 0.025 μg ml^−1^ penicillin; 100 μg ml^−1^ lysozyme; 0.025 μg/ml moenomycin, 500 μl DMSO, 0.05 μg ml^−1^ or 0.5 μg ml^−1^ tunicamycin and plates incubated at 37°C unless otherwise stated. Where indicated, plates were supplemented with 1 mM IPTG or 20 mM MgCl_2_. Images of plates were taken after 20-24 hours of incubation.

### 2.6 Cell length analysis

Overnight cultures of the indicated *L. monocytogenes* strains were inoculated to an OD_600_ of 0.1 and grown to an OD_600_ of 0.3-0.6 at 37°C with agitation in BHI medium or BHI medium containing 1 mM IPTG and the appropriate antibiotic. To stain the bacterial membranes, 800 μl of the bacterial cultures were mixed with 40 μl of 100 μg ml^−1^ nile red and incubated for 20 min at 37°C. The cells were washed twice in PBS buffer and subsequently re-suspended in the same buffer. Cells were then fixed in 1.12% paraformaldehyde for 20 min at room temperature in the dark and 1-1.5 μl of the cell suspension was spotted on microscope slides covered with a thin agarose layer (1.5% in ddH_2_O). Phase contrast and fluorescence images were taken using a Zeiss Axioskop 40 microscope equipped with an EC Plan-NEOFLUAR 100X/1.3 objective (Carl Zeiss, Göttingen, Germany) and coupled to an AxioCam MRm camera. Filter set 43 was used for the detection of the nile red signal. The images were processed using the Axio Vision software (release 4.7). The length of 50 cells per replicate was measured for each strain and the mean calculated. Statistical analysis was performed using the software GraphPad Prism (version 8).

### 2.7 Isolation of cellular proteins and western blotting

Bacteria from a 20 ml culture were harvested by centrifugation and washed with ZAP buffer (10 mM Tris-HCl, pH 7.5, 200 mM NaCl). The cells were subsequently resuspended in 1 ml ZAP buffer containing 1 mM PMSF and disrupted by sonication. Cellular debris was removed by centrifugation, the resulting supernatant collected and separated by SDS polyacrylamide gel electrophoresis (PAGE). Proteins were transferred onto positively charged polyvinylidene fluoride (PVDF) membranes using a semi-dry transfer unit. MurA was detected using a polyclonal rabbit antiserum raised against the *B. subtilis* MurAA protein (Kock et al., 2004), which also cross reacts with the *L. monocytogenes* MurA protein, as primary antibody and an anti-rabbit immunoglobulin G conjugated to horseradish peroxidase antibody as the secondary antibody. Blots were developed using ECL chemiluminescence reagents (Thermo Scientific) and imaged using a chemiluminescence imager (Vilber Lourmat).

### 2.8 Cytochrome C binding assay

Cytochrome C binding assays were performed as previously described (Kang et al., 2015). Briefly, overnight cultures of *L. monocytogenes* were used to inoculated fresh cultures to an OD_600_ of 0.05 in 4 ml BHI medium and the cultures were grown to an OD_600_ of 0.6-0.8 at 37°C and 200 rpm. Bacteria from 2 ml of the culture were harvested by centrifugation at 16.200 × g for 1 min. Cells were then washed twice with 20 mM MOPS (3-(N morpholino) propanesulfonic acid) buffer (pH 7) and adjusted to an OD_600_ of 0.25 in the same buffer. Cytochrome C was added at a final concentration of 50 μg ml^−1^ and the suspension was incubated in the dark for 10 min at room temperature. The suspension was centrifuged for 5 min at 16.200 × g, the supernatant removed and the absorbance of the supernatant was measured at 410 nm (OD_410_ + cells). A reaction without cells was used as blank (OD_410_ – cells). The percentage of bound cytochrome C was calculated as follows:

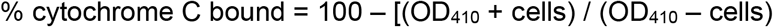

### 2.9 β-galactosidase assay

To compare the *dlt* promoter activity of the *L. monocytogenes* wildtype and suppressor strain ANG5746, β-galactosidase assays were performed. For this purpose, promoter-*lacZ* fusions were integrated into the *amyE* locus of *B. subtilis* 168. The resulting *B. subtilis* strains were grown in CSE-glucose minimal medium at 37°C to an OD_600_ of 0.5-0.8, bacteria from a culture aliquot harvested and the β-galactosidase activity determined as described previously (Miller, 1972).

## 3. RESULTS

### 3.1 Δ*eslB* phenotypes are restored in the presence of excess Mg^2+^

The *eslABCR* operon encodes the putative ABC transporter EslABC and the RpiR transcriptional regulator EslR. Absence of the transmembrane component EslB leads to a multitude of phenotypes including a cell division defect, decreased lysozyme resistance, the production of a thinner cell wall and a growth defect in media containing high sugar concentrations (Rismondo et al., 2021b). In addition, we observed that the growth of the *eslB* mutant is severely affected when grown on BHI plates at 42°C (Fig. 3A). So far, the cellular function of the transporter EslABC and how it is mechanistically linked to cell division and cell wall biosynthesis in *L. monocytogenes* is unknown. Generally, many cell wall defects can be rescued by the addition of Mg^2+^, however, the reason for this is still debated. Recently, it was speculated that binding of Mg^2+^ to the cell wall inhibits peptidoglycan hydrolases and thereby stabilizes the bacterial cell wall (Tesson et al., 2022). As the phenotypic defects of the *eslB* mutant suggest that peptidoglycan biosynthesis might be impaired, we speculated that the addition of Mg^2+^ should rescue the growth of this strain. To test this hypothesis, the *L. monocytogenes* wildtype 10403S, the Δ*eslB* mutant and the Δ*eslB* complementation strain were grown on BHI plates containing sucrose and penicillin, lysozyme or were incubated at 42°C in the absence or presence of 20 mM MgCl_2._ The addition of Mg^2+^ could restore the growth of the Δ*eslB* mutant under all conditions tested (Fig. S1), further supporting the hypothesis that peptidoglycan biosynthesis is impaired in *L. monocytogenes* cells lacking EslB.

**Figure 2:**
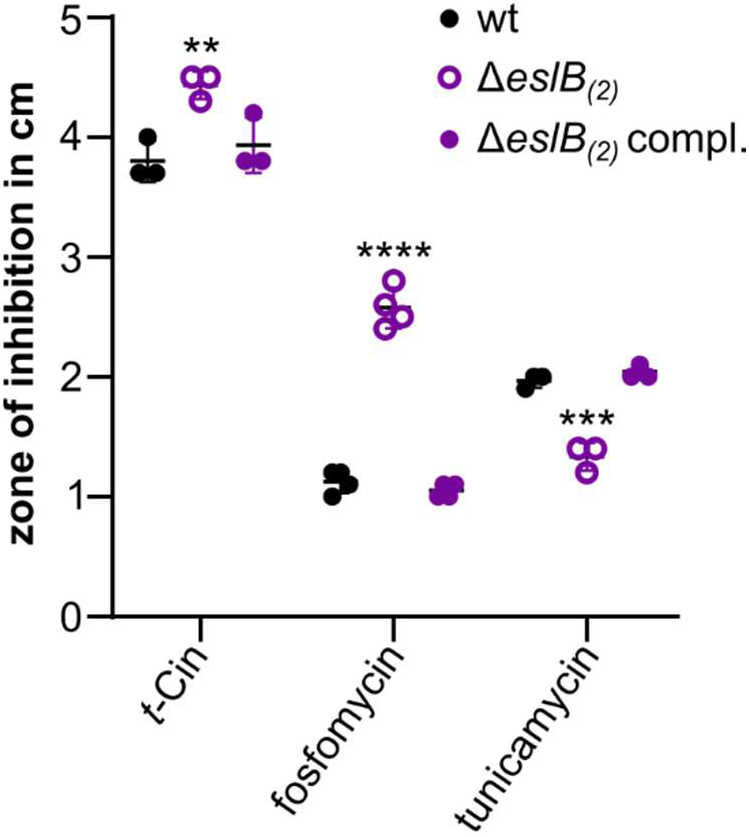
Alterations in the resistance of the *eslB* mutant towards cell wall-targeting antibiotics. Disk diffusion assay. *L. monocytogenes* strains 10403S (wt), Δ*eslB*_(*2*)_ and Δ*eslB*_(*2*)_ compl. were spotted on BHI plates. Antibiotic-soaked disks placed on top the agar surface and the plates incubated for 24h at 37°C. The inhibition zones for the indicated strains were measured and the average values and standard deviation of at least three independent experiments were plotted. For statistical analysis, a one-way ANOVA coupled with a Dunnett‘s multiple comparison test was used (** *p* ≤ 0.01, *** *p* ≤ 0.001, **** *p* ≤ 0.0001).

**Figure 3:**
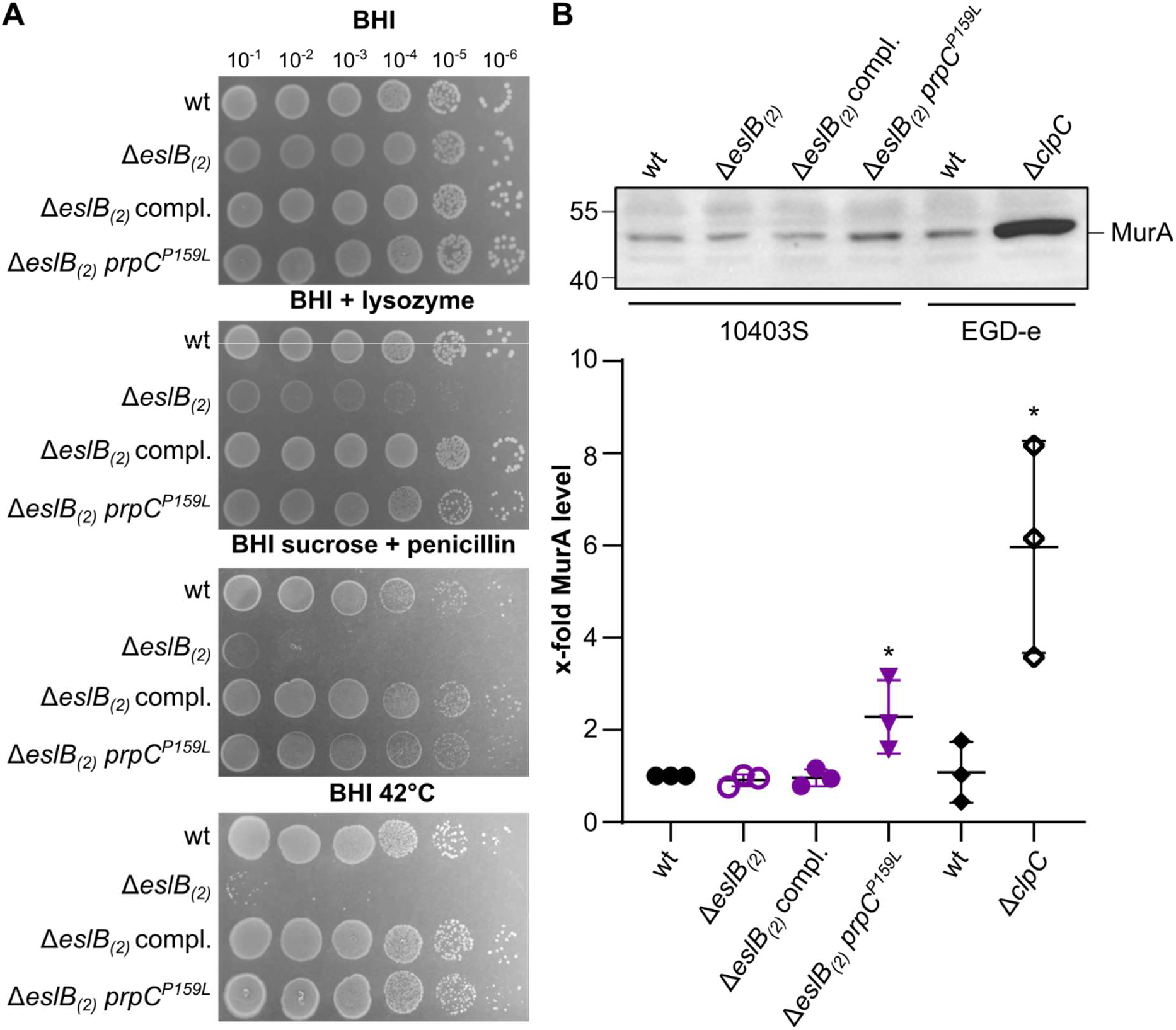
Mutation P159L in *prpC* suppresses *eslB* phenotypes. (A) Drop dilution assay. Dilutions of *L. monocytogenes* strains 10403S (wt), Δ*eslB*_(*2*)_, Δ*eslB*_(*2*)_ compl., and Δ*eslB*_(*2*)_ *prpC*^*P159L*^ were spotted on BHI plates, BHI plates containing 100 μg/ml lysozyme or containing 0.5 M sucrose and 0.025 μg/ml penicillin G and were incubated at 37°C or on BHI plates and were incubated at 42°C. (B) Western blot. Protein samples of strains 10403S, Δ*eslB*_(*2*)_, Δ*eslB*_(*2*)_ compl. and Δ*eslB*_(*2*)_ *prpC*^*P159L*^ were separated by SDS PAGE and transferred to a PVDF membrane. MurA was detected using a MurAA-specific antibody as described in the methods section. Protein samples of *L. monocytogenes* wildtype strain EGD-e and a *clpC* mutant were used as controls. MurA level in the protein samples were quantified and plotted. For statistical analysis, an unpaired two-tailed t-test was used (* *p* ≤ 0.05).

### 3.2 Resistance profile of the *eslB* mutant towards cell wall-targeting antibiotics

To narrow down which point of the peptidoglycan biosynthesis pathway is impaired in the *eslB* mutant, we performed disk diffusion assays with antibiotics that target different steps of this pathway (Fig. 1). Compared to the wildtype, the *eslB* mutant was more sensitive towards *t*-Cin and fosfomycin (Fig. 2), which target the UDP-Glc*N*Ac biosynthetic pathway and MurA, respectively (Marquardt et al., 1994; Pensinger et al., 2021; Sun et al., 2021). In contrast, the resistance towards D-cycloserine, nisin, vancomycin, moenomycin, penicillin, ampicillin and bacitracin (Fig. 1), which target processes downstream of MurA, was not altered in the *eslB* mutant (Fig. S2). The *eslB* mutant was more resistant towards tunicamycin, whose main target at low concentrations is TarO, the first enzyme functioning in the WTA biosynthesis pathway. As we have seen above, the inhibition of WTA biosynthesis seems to be beneficial for the *eslB* mutant likely due to an increased flux of UDP-Glc*N*Ac towards peptidoglycan biosynthesis. This antibiotic screen therefore suggests that one of the limiting factors of the *eslB* mutant might be the synthesis or correct distribution of UDP-Glc*N*Ac, a precursor, which is used for both peptidoglycan biosynthesis and the synthesis and modification of WTA. This hypothesis is supported by the observation that overproduction of GlmM and GlmR, two proteins involved in the production of UDP-Glc*N*Ac, can partially suppress the *eslB* phenotypes (Fig. S3).

### 3.3 Isolation of *eslB* suppressor mutants

To gain insights into the function of EslABC, we took advantage of the observation that the *eslB* mutant forms suppressors when grown on BHI plates that contain either 100 μg ml^−1^ lysozyme, or 0.5 M sucrose and 0.025 or 0.05 μg ml^−1^ penicillin. Genomic alterations present in independently isolated *eslB* suppressors were determined by whole genome sequencing. A large subset of *eslB* suppressors had mutations in *walR* (*lmo0287*) or *walK* (*lmo0288*), which encode the WalRK two-component system that is involved in cell wall metabolism (Dubrac et al., 2008; Howell et al., 2003). In addition, we identified mutations that mapped to genes associated with peptidoglycan biosynthesis (*murZ* (*lmo2552*), *reoM* (*lmo1503*), *prpC* (*lmo1821*), *pbpA1* (*lmo1892*)), peptidoglycan hydrolysis (*cwlO* (*lmo2505*, *spl*), *ftsX* (*lmo2506*), *ftsE* (*lmo2507*)) and wall teichoic acid (WTA) biosynthesis and modification (*tarL* (*lmo1077*), *dltX* (promoter region, *lmrg_02074*)) (Table 1).

**Table 1:**
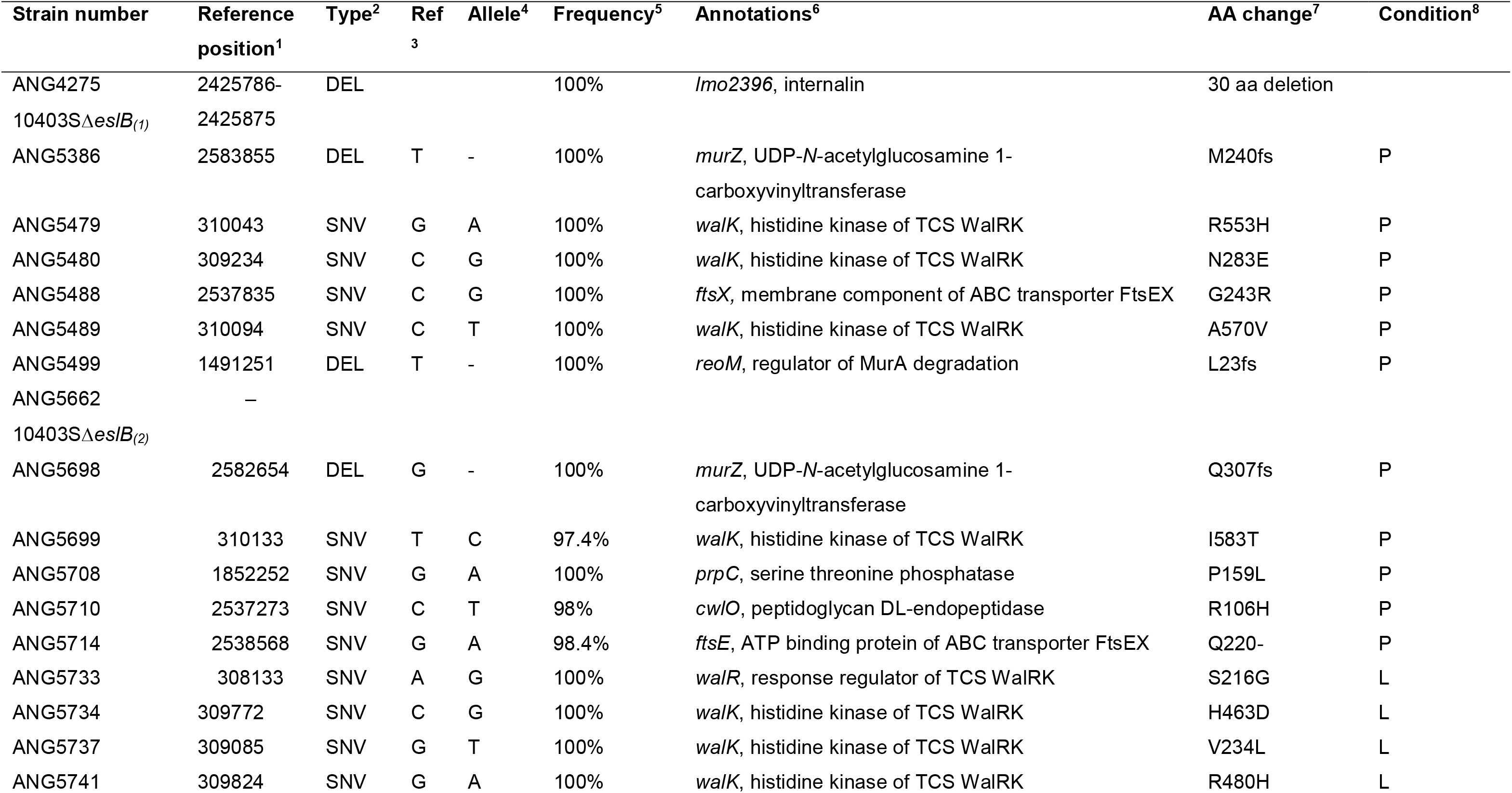
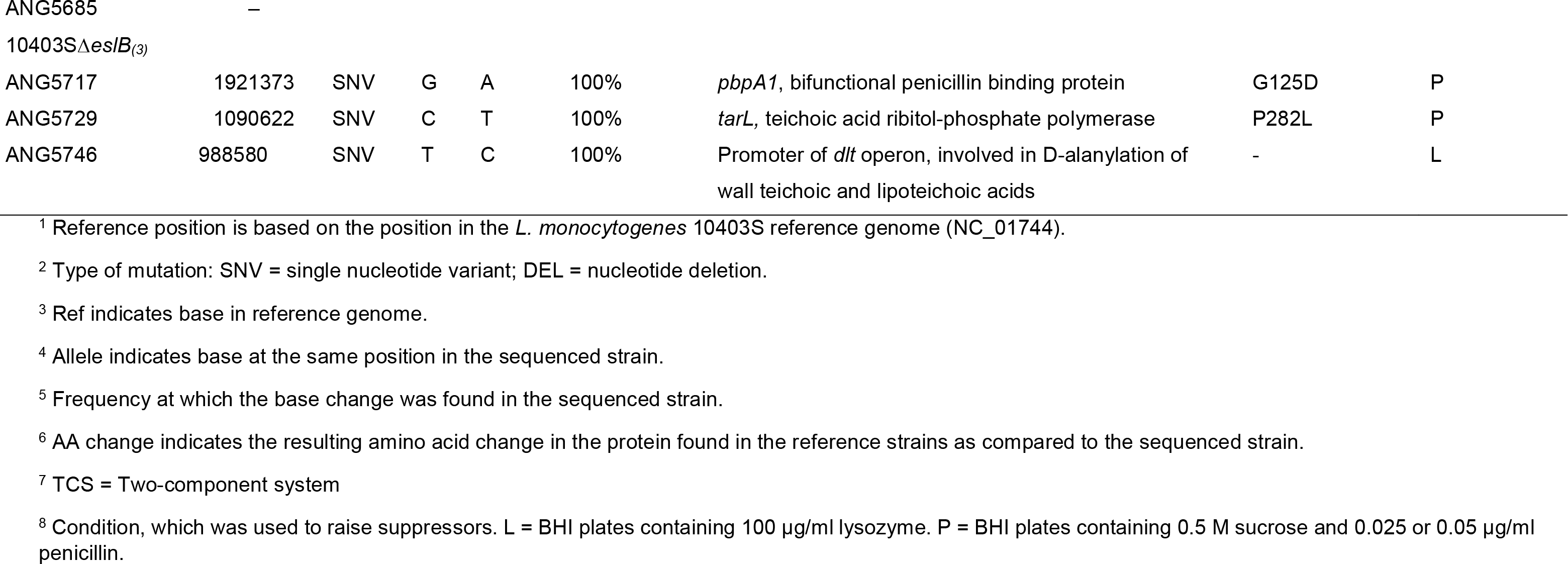
Identified sequence alterations in *L. monocytogenes eslB* deletion strains and suppressors.

### 3.4 Suppression of *eslB* phenotypes by increased MurA levels

Under sucrose penicillin stress, two *eslB* suppressors with mutations in *murZ* and one *eslB* suppressor with a mutation in *reoM* or *prpC* were isolated. The proteins encoded by all three of these genes affect MurA protein levels (Wamp et al., 2022, 2020). Drop dilution assays were performed to assess, whether mutations in *murZ*, *reoM* and *prpC* suppress the growth defect of the *eslB* mutant in the presence of sucrose and penicillin as well as other conditions under which the growth of the *eslB* mutant is severely affected. As expected, the growth of the *L. monocytogenes* strains 10403SΔ*eslB*_(*1*)_ *murZ*^*M240fs*^, 10403SΔ*eslB*_(*2*)_ *murZ*^*Q307fs*^, 10403SΔ*eslB*_(*1*)_ *reoM*^*K23fs*^ and 10403SΔ*eslB*_(*2*)_ *prpC*^*P159L*^ on BHI plates containing sucrose and penicillin is comparable to the wildtype strain 10403S. The suppressors with mutations in *murZ*, *reoM* and *prpC* also grew like wild-type on BHI plates containing lysozyme as well as on BHI plates that were incubated at 42°C (Fig. 3A, data not shown).

MurZ is a homolog of the UDP-*N*-acetylglucosamine 1-carboxyvinyltransferase MurA, which is required for the first step of peptidoglycan biosynthesis (Fig. 1)(Du et al., 2000; Kock et al., 2004). While MurA is essential for the growth of *L. monocytogenes*, the deletion of *murZ* is possible and results in the stabilization of MurA (Rismondo et al., 2016). MurA is a substrate of the ClpCP protease and a recent study showed that ReoM is required for the ClpCP-dependent proteolytic degradation of MurA in *L. monocytogenes* (Rismondo et al., 2016; Wamp et al., 2020). The activity of ReoM is controlled by the serine/threonine kinase PrkA and the cognate phosphatase PrpC. The phosphorylated form of ReoM stimulates peptidoglycan biosynthesis by preventing ClpCP-dependent degradation of MurA, while MurA degradation is enhanced in the presence of non-phosphorylated ReoM (Wamp et al., 2022, 2020). The identified mutations in *murZ* and *reoM* in our suppressor strains lead to frameshifts, and thus to the production of inactive MurZ and ReoM proteins. It has been previously shown that MurA protein levels are increased in *murZ* and *reoM* mutants (Rismondo et al., 2016; Wamp et al., 2020) and we thus assume that this is also the case for the *eslB* suppressor strains 10403SΔ*eslB*_(*1*)_ *murZ*^*M240fs*^, 10403SΔ*eslB*_(*2*)_ *murZ*^*Q307fs*^ and 10403SΔ*eslB*_(*1*)_ *reoM*^*K23fs*^. The identified mutation in *prpC* in the suppressor strain 10403SΔ*eslB*_(*2*)_ *prpC*^*P159L*^ leads to the production of a variant of PrpC, in which proline at position 159 is replaced by a leucine, however, it is currently not known whether this amino acid exchange leads to decreased or enhanced activity of PrpC. A decreased activity of PrpC would lead to accumulation of MurA (Wamp et al., 2022), while enhanced activity would result in reduced MurA levels. To assess which affect the P159L mutation in *prpC* has on MurA levels, western blots were performed using a MurAA-specific antibody. Protein samples of *L. monocytogenes* strains EGD-e and EGD-e Δ*clpC* were used as controls. In accordance with previous studies, MurA accumulated in the *clpC* mutant (Rismondo et al., 2016). The production of the mutated PrpC variant, PrpC^P159L^, also led to a slight accumulation of MurA in the *eslB* mutant background (Fig. 3B). Additionally, no change in MurA levels could be observed for the *eslB* mutant compared to the wildtype strain 10403S (Fig 3B). These results suggest that while the decrease in peptidoglycan production seen in the *eslB* mutant is not caused by decreased MurA levels, the growth deficit of the *eslB* mutant can be rescued by preventing MurA degradation.

To confirm that the suppression of the *eslB* phenotypes in the strains with mutations in *murZ*, *reoM* or *prpC* is indeed the result of increased MurA protein levels, we integrated a second, IPTG-inducible copy of *murA* into the genome of the *eslB* mutant. First, we determined the resistance towards fosfomycin, which is a known inhibitor of MurA (Fig. 4A-B)(Kahan et al., 1974), for the *L. monocytogenes* wildtype and the *eslB* mutant harboring the empty plasmid pIMK3 or pIMK3-*murA* in the presence of IPTG. Strain 10403SΔ*eslB* pIMK3 is three-fold more sensitive to fosfomycin as compared to the cognate wildtype strain. The induction of *murA* expression increases the resistance of strains 10403S pIMK3-*murA* and 10403SΔ*eslB* pIMK3-*murA* (Fig. 4A-B), suggesting that MurA is indeed overproduced in these strains.

**Figure 4:**
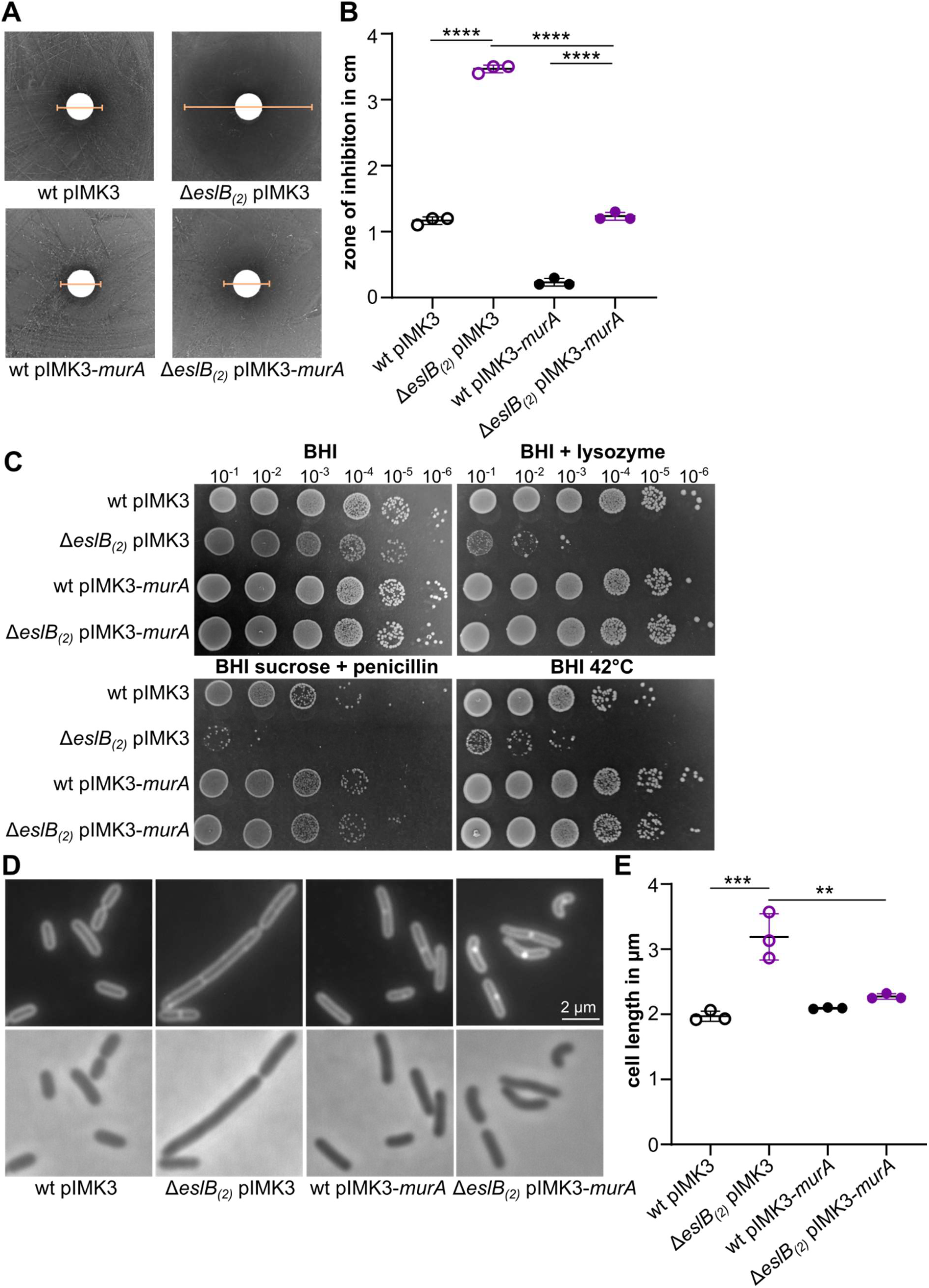
MurA overexpression leads to suppression of *eslB* phenotypes. (A-B) Fosfomycin disk diffusion assay. (A) *L. monocytogenes* strains wt pIMK3, Δ*eslB* pIMK3, wt pIMK3-*murA* and Δ*eslB* pIMK3-*murA* were plated on BHI plates containing 1 mM IPTG. Fosfomycin-soaked disks were placed on the agar surface and the plates incubated for 24 h at 37°C. Yellow lines indicate the diameter of the zone of inhibition. (B) The inhibition zones for the indicated strains were measured and the average values and standard deviation of three independent experiments were plotted. (C) Drop dilution assay. Dilutions of *L. monocytogenes* strains 10403S pIMK3 (wt pIMK3), Δ*eslB* pIMK3, wt pIMK3-*murA* and Δ*eslB* pIMK3-*murA* were spotted on BHI plates, BHI plates containing 100 μg/ml lysozyme or containing 0.5 M sucrose and 0.025 μg/ml penicillin G, which were incubated at 37°C or on BHI plates, which were incubated at 42°C. (D) Microscopy images of *L. monocytogenes* strains. Bacterial membranes were stained with nile red as described in the methods section. Scale bar is 2 μm. (E) Cell length of *L. monocytogenes* strains shown in panel B. The cell length of 50 cells per strain was determined and the median cell length calculated. The average values and standard deviations of three independent experiments are plotted. For statistical analysis, a one-way ANOVA coupled with Tukey‘s multiple comparison test was used (** *p* ≤ 0.01, *** *p* ≤ 0.001, **** *p* ≤ 0.0001).

Next, we assessed whether the growth phenotypes of the *eslB* mutant can be suppressed by the overexpression of *murA*. In the presence of the inducer IPTG, the growth of strain 10403SΔ*eslB* pIMK3-*murA* was comparable to the corresponding wildtype strain under all conditions tested (Fig. 4C). These results indicate that overproduction of MurA can compensate for the loss of *eslB* and that the suppression of the *eslB* phenotypes in the *murZ*, *reoM* and *prpC* suppressors is likely the result of increased MurA levels.

Interestingly, increased MurA production also leads to the suppression of the cell division defect of the *eslB* mutant. Cells of the *eslB* mutant carrying the empty pIMK3 plasmid have a cell length of 3.19±0.29 μm. In contrast, strain 10403SΔ*eslB* pIMK3-*murA* produces cells with a length of 2.27±0.03 μm, a size that is comparable to the length of *L. monocytogenes* wildtype cells (Fig. 4D-E).

### 3.5 Suppression by reduction of the glycosyltransferase activity of PBP A1

Multiple enzymes are involved in the production of lipid II, the peptidoglycan precursor, within the cytoplasm. After its transport across the membrane via MurJ, lipid II is incorporated into the growing glycan strand by glycosyltransferases, which can either be class A penicillin binding proteins (PBPs), monofunctional glycosyltransferases or members of the SEDS (shape, elongation, division, sporulation) family, such as RodA and FtsW (Cho et al., 2016; Emami et al., 2017; Meeske et al., 2016). *L. monocytogenes* encodes two class A PBPs, PBP A1 and PBP A2, in its genome (Korsak et al., 2010; Rismondo et al., 2015). One of the *eslB* suppressor strains, ANG5717, carries a point mutation in *pbpA1* leading to the substitution of glycine at position 125 by aspartate. Interestingly, this glycine residue is part of the catalytic site of the glycosyltransferase domain, suggesting that the glycosyltransferase activity of the PBP A1 variant PBP A1^G125D^ might be reduced. Drop dilution assays and microscopic analysis revealed that the phenotypes associated with deletion of *eslB* can be rescued by *pbpA1*^*G125D*^ (Fig. 5A-C). The glycosyltransferase activity of class A PBPs can be inhibited by moenomycin (Huber and Nesemann, 1968). We thus tested whether the presence of moenomycin would enable the growth of the *eslB* mutant under heat stress. As shown in Figure 5D, the *eslB* mutant is unable to grow at 42°C on BHI plates or BHI plates containing DMSO, which was used to dissolve moenomycin. In contrast, nearly wildtype-like growth of the *eslB* mutant was observed on BHI plates that were supplemented with moenomycin. These results suggest that growth deficits of the *eslB* mutant can be rescued by the reduction of the glycosyltransferase activity of class A PBPs.

**Figure 5:**
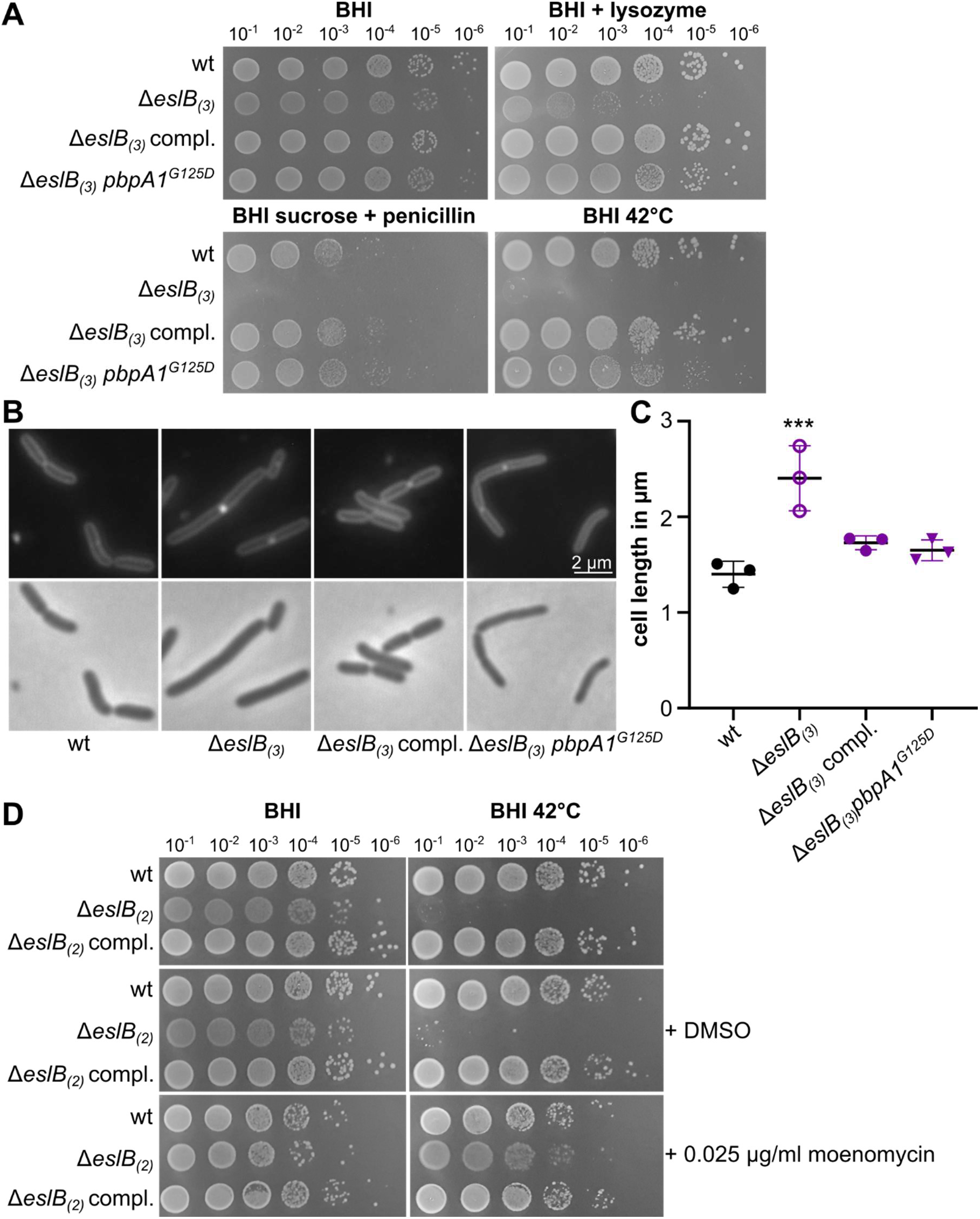
Inactivation of the glycosyltransferase of penicillin binding proteins rescues the *eslB* mutant. (A) Drop dilution assay. Dilutions of *L. monocytogenes* strains 10403S (wt), Δ*eslB*_(*3*)_, Δ*eslB*_(*3*)_ compl. and Δ*eslB*_(*3*)_ *pbpA1*^*G125D*^ were spotted on BHI plates, BHI plates containing 100 μg/ml lysozyme or containing 0.5 M sucrose and 0.025 μg/ml penicillin G, which were incubated at 37°C or on BHI plates, which were incubated at 42°C. (B) Microscopy images of *L. monocytogenes* strains. Bacterial membranes were stained with nile red as described in the methods section. Scale bar is 2 μm. (C) Cell length of *L. monocytogenes* strains shown in panel B. The cell length of 50 cells per strain was determined and the median cell length calculated. The average values and standard deviations of three independent experiments are plotted. (D) Drop dilution assay. Dilutions of *L. monocytogenes* strains 10403S (wt), Δ*eslB*_(*2*)_ and Δ*eslB*_(*2*)_ compl. were spotted on BHI plates, BHI plates containing DMSO or containing 0.025 μg/ml moenomycin and incubated at 37°C or 42°C.

### 3.6 Suppression by reducing *cwlO* expression or CwlO activity

In addition to suppressors associated with MurA and PBP A1, we isolated suppressors carrying mutations in genes that either affect the transcription of *cwlO* or the activity of CwlO. CwlO is a DL-endopeptidase, which opens the existing cell wall to allow for the insertion of new lipid II precursors during cell elongation of *B. subtilis* (Bisicchia et al., 2007; Hashimoto et al., 2012). In *B. subtilis*, transcription of *cwlO* is induced by WalRK in response to low DL-endopeptidase activity (Bisicchia et al., 2007; Dobihal et al., 2019; Dubrac et al., 2008) and CwlO activity is stimulated by a direct interaction with the ABC transporter FtsEX (Meisner et al., 2013). In our suppressor screen, we isolated eight *eslB* suppressor strains that carried mutations in *walK*, which were either isolated under lysozyme or sucrose penicillin stress, and four *eslB* suppressor strains that carried mutations in *walR* from which the majority was isolated in the presence of lysozyme stress. We selected *eslB* suppressors 10403SΔ*eslB*_(*2*)_ *walR*^*S216G*^ and10403SΔ*eslB*_(*2*)_ *walK*^*H463D*^, which were isolated under lysozyme stress, and 10403SΔ*eslB*_(*2*)_ *walK*^*I583T*^, which was isolated under sucrose penicillin stress, for further analysis. In addition, three suppressors with mutations were isolated in *ftsE*, *ftsX* and *cwlO* under sucrose penicillin pressure (Table 1).

Spot plating assays showed that the *eslB* suppressors with mutations in either *walR* or *walK* could overcome the growth defect of the *eslB* mutant under sucrose penicillin, lysozyme and heat stress (Fig. 6). In contrast, *eslB* suppressors harboring mutations in either *ftsE*, *ftsX* or *cwlO*, grow comparable to the wildtype under sucrose penicillin and heat stress, but showed a similar growth defect as the *eslB* mutant on plates containing lysozyme (Fig. 6A). Microscopic analysis of the *eslB* suppressors with mutations in *walR*, *walK*, *ftsE*, *ftsX* and *cwlO* revealed that these cells have a similar cell length to the *L. monocytogenes* wildtype 10403S and thus, the expression of mutated WalR, WalK, FtsE and CwlO variants could restore the cell division defect of the *eslB* mutant (Fig. 6B-C). Additionally, we observed that the cells of suppressor strains 10403SΔ*eslB*_(*2*)_ *cwlO*^*R106H*^ and 10403SΔ*eslB*_(*2*)_ *ftsE*^*Q220-*^ were bent compared to the wildtype strain. This phenotype is characteristic for *B. subtilis cwlO* and *ftsE* mutants (Domínguez-Cuevas et al., 2013; Meisner et al., 2013) suggesting that the acquired mutations in *cwlO* and *ftsE* led to inactivation or reduced activity of CwlO and FtsE in *L. monocytogenes*.

**Figure 6:**
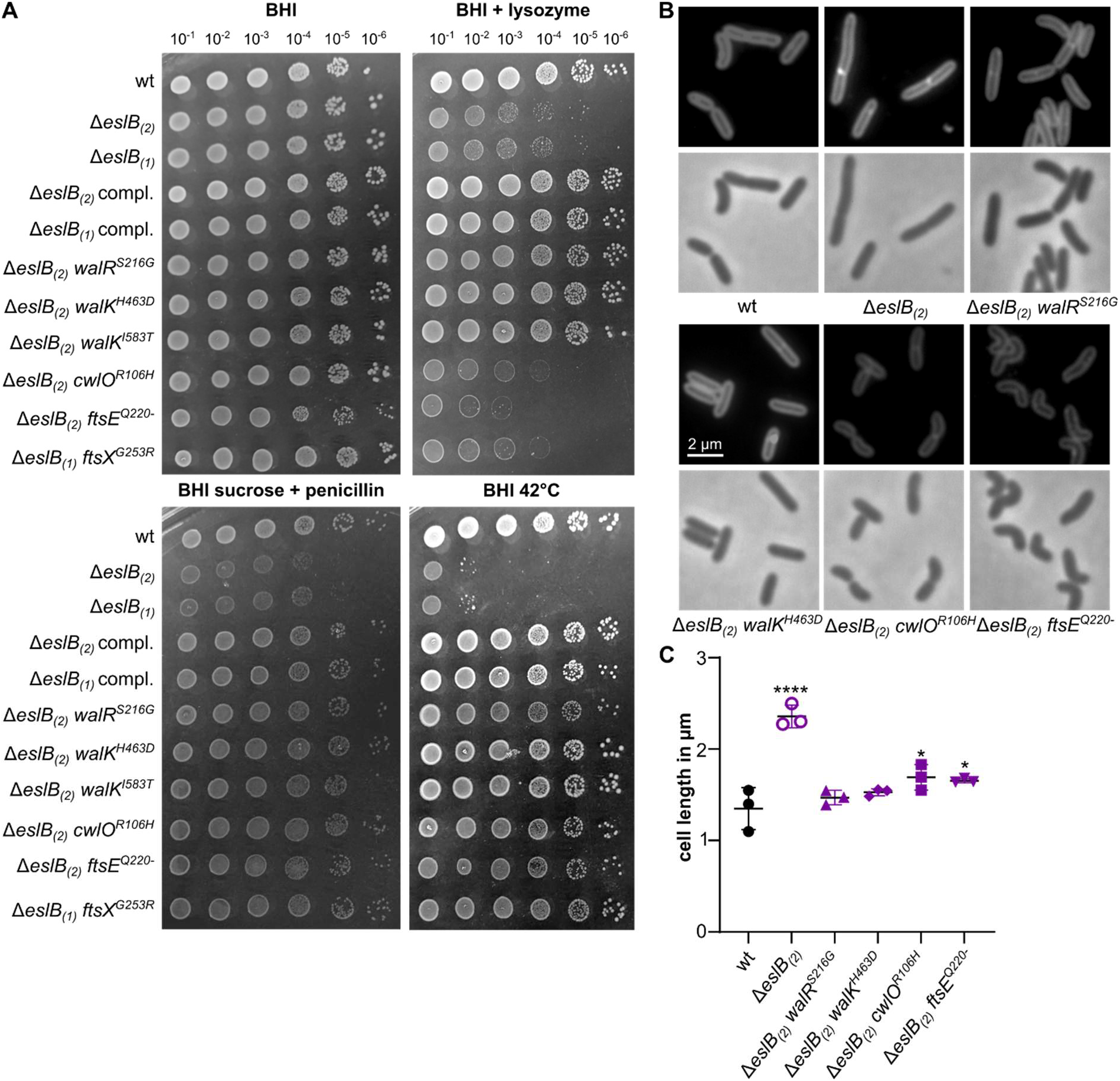
Mutations in *walR*, *walK*, *cwlO*, *ftsE* and *ftsX* suppress *eslB* phenotypes. (A) Drop dilution assay. Dilutions of *L. monocytogenes* strains 10403S (wt), Δ*eslB*_(*2*)_, Δ*eslB*_(*1*)_, Δ*eslB*_(*2*)_ compl., Δ*eslB*_(*1*)_ compl., Δ*eslB*_(*2*)_ *walR*^*S216G*^, Δ*eslB*_(*2*)_ *walK*^*H463D*^, Δ*eslB*_(*2*)_ *walK*^*I583T*^, Δ*eslB*_(*2*)_ *cwlO*^*R106H*^, Δ*eslB*_(*2*)_ f*tsE*^*Q220*−^ and Δ*eslB*_(*1*)_ *ftsX*^*G253R*^ were spotted on BHI plates, BHI plates containing 100 μg/ml lysozyme or containing 0.5 M sucrose and 0.025 μg/ml penicillin G and were incubated at 37°C or on BHI plates that were incubated at 42°C. (B) Microscopy images of *L. monocytogenes* strains. Bacterial membranes were stained with nile red as described in the methods section. Scale bar is 2 μm. (C) Cell length of *L. Monocytogenes* strains shown in panel B. The cell length of 50 cells per strain was determined and the median cell length calculated. The average values and standard deviations of three independent experiments are plotted. For statistical analysis, a one-way ANOVA coupled with a Dunnett‘s multiple comparison test was used (* *p* ≤ 0.05, **** *p* ≤ 0.0001).

The observation that mutations accumulate in genes associated with CwlO led to the hypothesis that wildtype CwlO activity is toxic, due to the reduced peptidoglycan levels produced by the *eslB* mutant (Rismondo et al., 2021b). To confirm this hypothesis, an *eslB cwlO* double mutant was constructed, in which the expression of an ectopic copy of *cwlO* was placed under the control of an IPTG-inducible promoter. As expected, in the absence of the inducer, the *eslB cwlO* double mutant was able to grow on BHI plates supplemented with sucrose and penicillin. In accordance to the results obtained for the suppressors, the growth of a strain lacking both, EslB and CwlO, was still impaired in the presence of lysozyme. Furthermore, individual deletion of either *eslB* or *cwlO* resulted in a severe growth defect at higher temperatures, while the *eslB cwlO* double mutant was able to grow in the absence of IPTG at 42°C. Induction of *cwlO* expression in strain 10403SΔ*cwlO* Δ*eslB* pIMK3-*cwlO* resulted in a growth defect under sucrose penicillin and under heat stress, which is comparable to that of the *eslB* mutant (Fig. 7A). Next, we studied the cell morphology of strains lacking CwlO, EslB or both. In accordance with previous studies investigating the function of CwlO in *B. subtilis* (Domínguez-Cuevas et al., 2013; Meisner et al., 2013), the *L. moncytogenes cwlO* mutant formed slightly smaller, bent cells (Fig.7B-C). Cells lacking both, EslB and CwlO, were also shorter than cells of the *L. monocytogenes* wildtype 10403S, while induction of *cwlO* expression in strain 10403SΔ*cwlO* Δ*eslB* pIMK3-*cwlO* resulted in the formation of elongated cells (Fig. 7B-C). Altogether, these findings support our hypothesis that the suppression of the *eslB* phenotypes in the *cwlO*, *ftsE* and *ftsX* suppressors is the result of an inactivation of the DL-endopeptidase CwlO.

**Figure 7:**
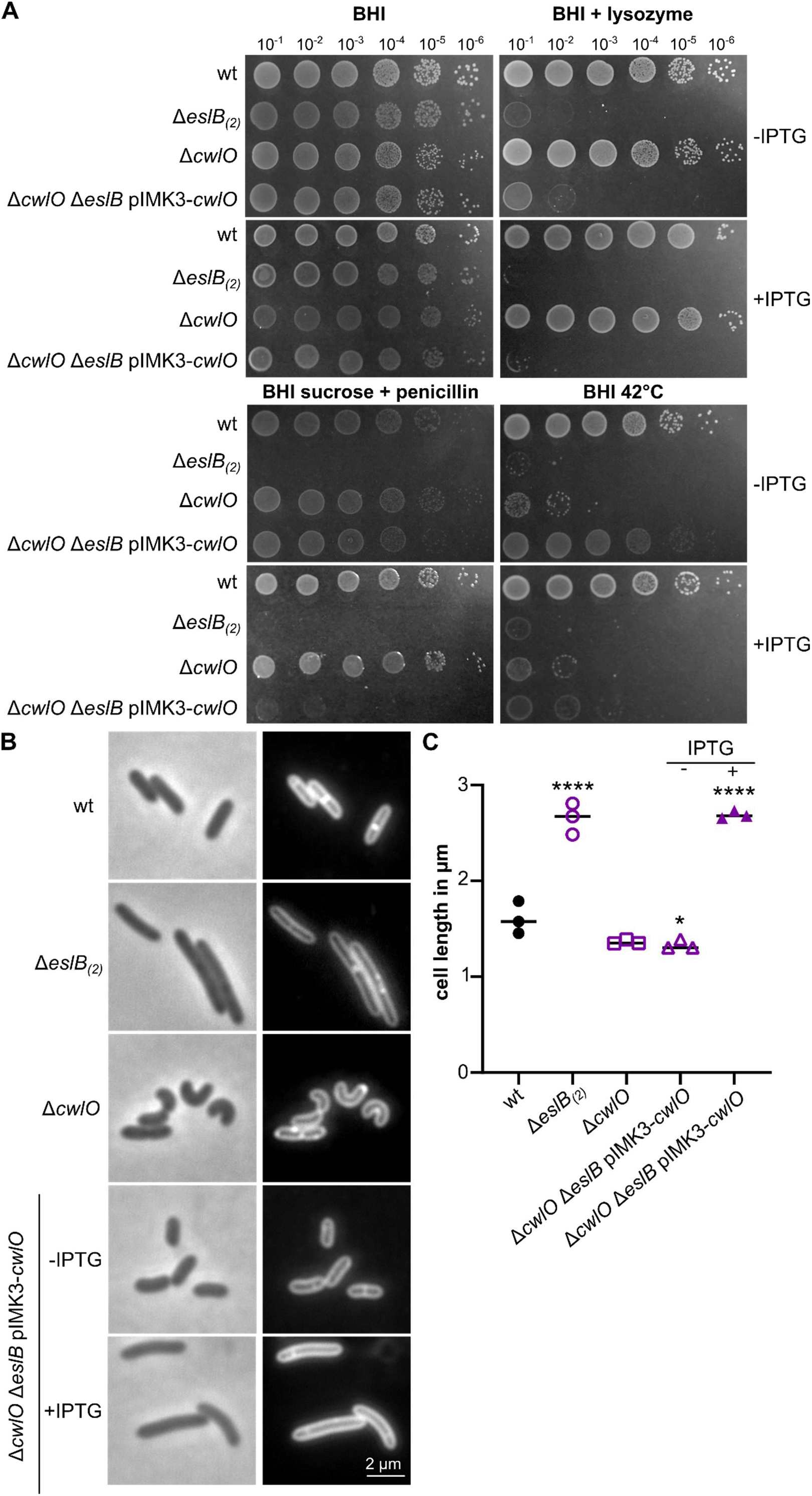
Absence of CwlO suppresses most of the *eslB* phenotypes. (A) Drop dilution assay. Dilutions of *L. monocytogenes* strains 10403S (wt), Δ*eslB*_(*2*)_, Δ*cwlO* and Δc*wlO* Δ*eslB* pIMK3-*cwlO* were spotted on BHI plates, BHI plates containing 100 μg/ml lysozyme or containing 0.5 M sucrose and 0.025 μg/ml penicillin G and were incubated at 37°C or on BHI plates that were incubated at 42°C. For induction of *cwlO* expression, the indicated plates were supplemented with 1 mM IPTG. (B) Microscopy images of *L. monocytogenes* strains. Bacterial membranes were stained with nile red as described in the methods section. Scale bar is 2 μm. (C) Cell length of *L. monocytogenes* strains shown in panel B. The cell length of 50 cells per strain was determined and the median cell length calculated. The average values and standard deviations of three independent experiments are plotted. For statistical analysis, a one-way ANOVA coupled with a Dunnett‘s multiple comparison test was used (* *p* ≤ 0.05, **** *p* ≤ 0.0001).

### 3.7 Suppression by inhibition of WTA biosynthesis

In *B. subtilis*, WalRK does not only stimulate transcription of *cwlO*, but also of the *tagAB* and *tagDEFGH* operons, which encode proteins required for WTA biosynthesis and export (Howell et al., 2003). In *L. monocytogenes* 10403S, WTA is composed of a ribitol phosphate backbone, which is attached to a Glc-Glc-glycerol phosphate-Glc*N*Ac-Man*N*Ac linker unit and modified with Glc*N*Ac, rhamnose and d-alanine residues (Brown et al., 2013; Shen et al., 2017). Thus, UDP-Glc*N*Ac is consumed by several enzymes during WTA biosynthesis and modification. As mentioned above, we isolated suppressors containing mutations in *lmo1077*, encoding a TarL homolog, and the promoter region of the *dlt* operon (Table 1). TarL is a teichoic acid ribitol phosphate polymerase required for the polymerization of the ribitol phosphate backbone of WTA (Fig. 1)(Brown et al., 2013). The *dlt* operon codes for proteins that are essential for the d-alanylation of WTA (Neuhaus and Baddiley, 2003; Perego et al., 1995; Rismondo et al., 2021a). Figure 8A shows that the *eslB* mutant, which harbors a mutation in *tarL*, could grow again under sucrose penicillin and lysozyme stress, likely due to a reduction in WTA biosynthesis and potentially leading to an increase in the cellular UDP-Glc*N*Ac pool, which can then be used by MurA. However, the *tarL* suppressor is unable to grow at 42°C. A similar phenotype could be observed when the *L. monocytogenes* wildtype strain 10403S is grown at 42°C on a plate containing high concentrations of tunicamycin, which inhibits TarO, the first enzyme required for WTA biosynthesis (data not shown). This indicates that the mutated TarL variant produced by the *eslB* suppressor strain ANG5729 has a reduced activity. To further test if inhibition of WTA biosynthesis is indeed beneficial for the *eslB* mutant, we grew the *eslB* mutant at 42°C on BHI plates containing different concentrations of tunicamycin. We could see partial suppression of the heat sensitivity at a tunicamycin concentration of 0.05 μg/ml and the *eslB* mutant grew comparable to the *L. monocytogenes* wildtype strain 10403S and the complementation strain on plates containing 0.5 μg/ml tunicamycin (Fig. S4). These results demonstrate that reduction of WTA biosynthesis leads to the suppression of the heat phenotype associated of the *eslB* mutant.

**Figure 8:**
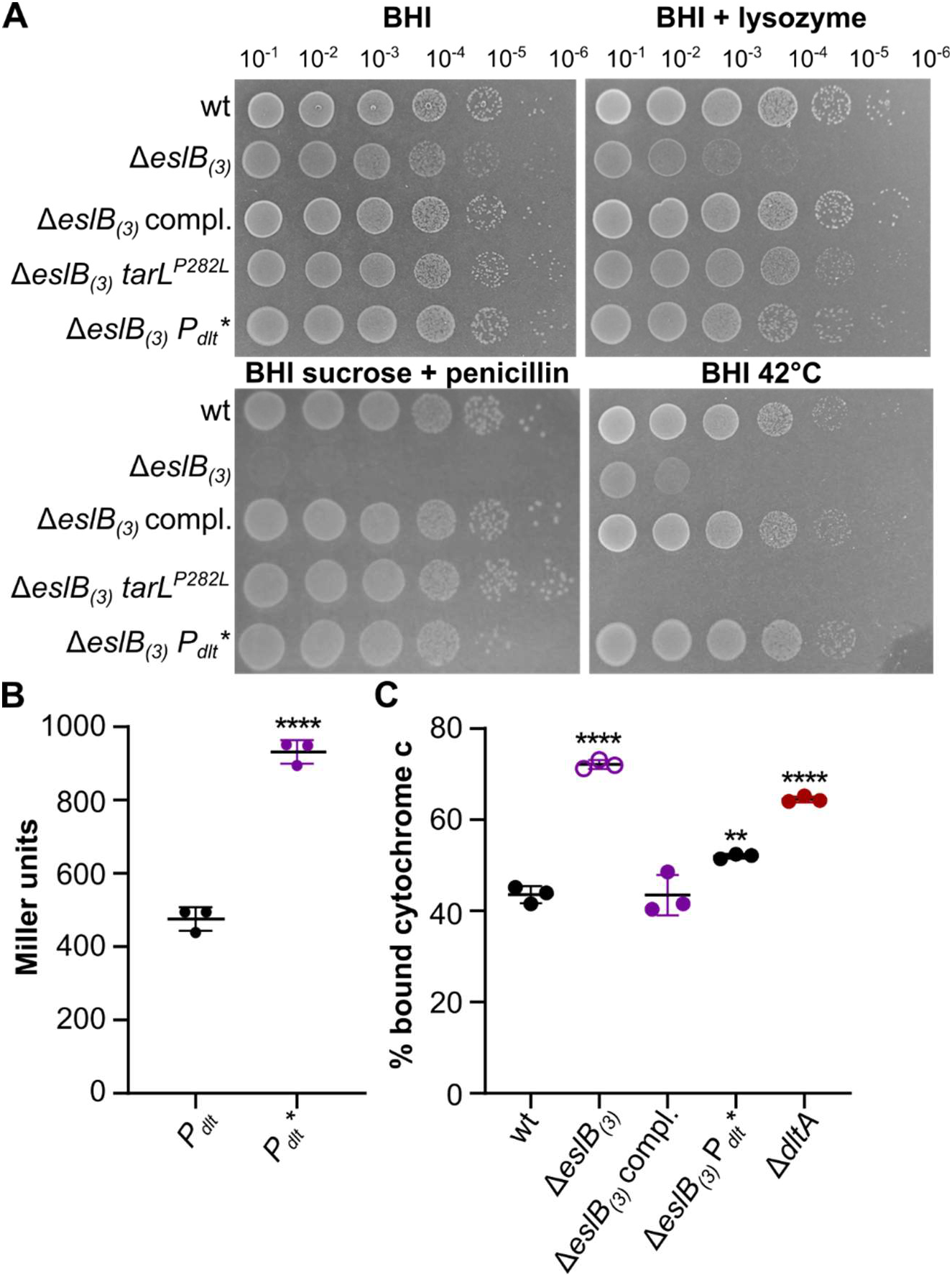
Suppression of *eslB* phenotypes by alterations in WTA biosynthesis and modification. (A) Drop dilution assay. Dilutions of *L. monocytogenes* strains 10403S (wt), Δ*eslB*_(*3*)_, Δ*eslB*_(*3*)_ compl., Δ*eslB*_(*3*)_ *tarL*^*P282L*^ and Δ*eslB*_(*3*)_ *P*_*dlt*_* were spotted on BHI plates, BHI plates containing 100 μg/ml lysozyme or containing 0.5 M sucrose and 0.025 μg/ml penicillin G and were incubated at 37°C or on BHI plates that were incubated at 42°C. (B) β-galactosidase assay. The *dlt* promoter of *L. monocytogenes* wildtype 10403S (*P*_*dlt*_) and suppressor strain Δ*eslB*_(*3*)_ *P*_*dlt*_* (*P*_*dlt*_*) were fused with *lacZ* and integrated into the *amyE* locus of *Bacillus subtilis*. The promoter activity was determined in Miller units as described in the methods section and the average and standard deviation of three independent experiments were plotted. An unpaired t-test was used for statistical analysis (**** *p* ≤ 0.0001). (C) Cytochrome C assay. Cells of *L. monocytogenes* strains 10403S, Δ*eslB*_(*3*)_, Δ*eslB*_(*3*)_ compl. and Δ*eslB*_(*3*)_ *P*_*dlt*_* were incubated with cytochrome C as described in the methods section. The percentage of cytochrome C, which is bound by the cell surface, was calculated for three independent experiments and plotted. A strain lacking DltA was used as control. For statistical analysis, a one-way ANOVA coupled with a Dunnett‘s multiple comparison test was used (** *p* ≤ 0.01, **** *p* ≤ 0.0001).

Figure 8A further shows that the *eslB* suppressor strain, which harbors a mutation in the *dlt* promoter region, could grow again under all conditions tested. To assess what consequence the point mutation in the promoter region of the *dlt* operon has on gene expression, we place the promoterless *lacZ* gene under the control of the *dlt* promoter or the mutated *dlt* promoter found in suppressor strain ANG5746. The *lacZ* promotor gene fusions were introduced into *B. subtilis* and β-galactosidase activities determined. This analysis showed that the mutation in the *dlt* promoter resulted in increased promoter activity and likely increased d-alanylation of teichoic acids in the suppressor strain (Fig. 8B). The d-alanylation state of teichoic acids is an important factor that impacts the bacterial cell surface charge (Vadyvaloo et al., 2004). As we isolated a suppressor that increased the expression of the *dlt* operon, we wondered, whether the surface charge of *eslB* mutant cells is altered. To test this, we determined the binding capability of positively charged cytochrome C to the cell surface of different *L. monocytogenes* strains, which serves as a readout of the bacterial surface charge (Peschel et al., 1999; Wecke et al., 1997). A strain lacking d-alanine residues on LTA and WTA due to a deletion of *dltA* was used as a control. As expected, the cell surface of the *dltA* mutant had a higher negative charge as compared to the *L. monocytogenes* wildtype strain 10403S. A similar result was observed for the *eslB* mutant. The high negative surface charge of the *eslB* mutant could be partially rescued by the mutation in the *dlt* promoter, which is present in the *eslB* suppressor strain 10403SΔ*eslB* P_*dlt*_* (Fig. 8C). This result suggests that either the d-alanylation state of WTAs is altered in the absence of EslB or that the surface presentation of WTA is changed due to the production of a thinner peptidoglycan layer.

## 4. DISCUSSION

In this study, we aimed to provide further insight into the connection between the predicted ABC transporter EslABC and peptidoglycan biosynthesis. Previous studies have shown that the deletion of *eslB*, coding for one of transmembrane components of the transporter, leads to a growth defect on sucrose containing media, the formation of elongated cells, the production of a thinner peptidoglycan layer, as well as sensitivity towards the natural antibiotic lysozyme and cationic antimicrobial peptides (Burke et al., 2014; Durack et al., 2015; Rismondo et al., 2021b), suggesting its involvement in peptidoglycan biosynthesis and cell division. It was proposed that the addition of Mg^2+^ rescues mutants with a defect in peptidoglycan biosynthesis by reducing the activity of peptidoglycan hydrolases (Tesson et al., 2022). In accordance with this, we observed a suppression of the *eslB* growth deficits under all conditions tested when an excess of Mg^2+^ was added. *L. monocytogenes* encodes several peptidoglycan hydrolases, including the DL-endopeptidase CwlO (Spl), the LytM-domain containing protein Lmo2504 and the two cell wall hydrolases NamA and CwhA (p60), which are required for daughter cell separation (Carroll et al., 2003; Pilgrim et al., 2003). We have isolated several *eslB* suppressors, which carry mutations in *cwlO* or mutations in other genes, e.g. *ftsE* or *ftsX*, which affect CwlO activity. After depletion of CwlO, the *eslB* mutant was able to grow in otherwise non-permissive conditions, suggesting that the activity of CwlO might be deregulated in the *eslB* deletion strain or that the peptidoglycan of the *eslB* mutant is more sensitive to hydrolysis by CwlO. CwlO activity is controlled by a direct protein-protein interaction with the ABC transporter FtsEX (Meisner et al., 2013). It is thus tempting to speculate that the ABC transporter EslABC might also affect CwlO activity or localization of proteins involved in peptidoglycan biosynthesis and/or degradation. Interestingly, a strain lacking both EslB and CwlO is able to grow at elevated temperatures, while both single mutants are not able to grow under this condition.

We could isolate *eslB* suppressor strains with mutations that are associated with stabilization of MurA, the UDP-Glc*N*Ac 1-carboxyvinyltransferase responsible for the first step of peptidoglycan biosynthesis. Elevated levels of MurA, and thus, enhanced peptidoglycan biosynthesis could fully restore the phenotypic defects of the 10403SΔ*eslB* strain. To identify the exact process of peptidoglycan biosynthesis, which is impaired in the *eslB* mutant, we performed a screen with antibiotics targeting different steps of this process. The absence of *eslB* only affected the resistance towards *t*-Cin and fosfomycin, which reduce the synthesis of UDP-Glc*N*Ac and inhibit MurA, respectively (Marquardt et al., 1994; Pensinger et al., 2021; Sun et al., 2021). Surprisingly, we observed an increased resistance of the *eslB* mutant towards tunicamycin. The primary target of tunicamycin is TarO, the first enzyme of WTA biosynthesis, and, at high concentrations, also MraY, which is responsible for the production of lipid I during peptidoglycan biosynthesis (Fig. 1)(Campbell et al., 2011; Hakulinen et al., 2017; Price and Tsvetanova, 2007; Watkinson et al., 1971). During WTA biosynthesis, UDP-Glc*N*Ac is used for the synthesis of the linker unit and for the modification of the WTA backbone (Eugster et al., 2015; Rismondo et al., 2018; Shen et al., 2017). Thus, reducing the production of WTA could increase the availability of UDP-Glc*N*Ac for peptidoglycan biosynthesis and therefore support the growth of the *eslB* mutant under non-permissive conditions. In accordance with this, we observed a partial suppression of the *eslB* phenotypes by the overproduction of GlmM and GlmR, two enzymes required for the synthesis of UDP-Glc*N*Ac (Pensinger et al., 2021). These results suggest that either the production or distribution of UDP-Glc*N*Ac as a substrate between different pathways might be disturbed in the *eslB* mutant.

Cell wall stability depends on the balance between peptidoglycan biosynthesis and hydrolysis. An imbalance of one of these processes results in rapid cell lysis (Sassine et al., 2020). Recent studies suggest that new peptidoglycan precursors are inserted into the growing glycan chains by the Rod system, which is composed of several enzymes. In contrast, class A PBPs are thought to fill gaps and/or repair cell wall defects (Cho et al., 2016; Dion et al., 2019; Vigouroux et al., 2020). It was recently shown that enhanced endopeptidase activity leads to the activation of class A PBPs in *E. coli* (Lai et al., 2017). A similar mechanism seems to exist in *B. subtilis*, as either inactivation of PBP1 or inhibition of peptidoglycan hydrolases by the addition of Mg^2+^ suppresses growth and morphological defects of an *mreB* mutant (Tesson et al., 2022). In accordance with this, we also observed suppression of the heat sensitivity of the *eslB* mutant in presence of moenomycin, which specifically inhibits the glycosyltransferase activity of class A PBPs (Ostash and Walker, 2010; Van Heijenoort et al., 1978).

In our suppressor screen, we identified a strain carrying a mutation in the *dlt* promoter region, which leads to an overproduction of the Dlt enzymes. These enzymes are required for the modification of teichoic acids (TAs) with d-alanines (Neuhaus and Baddiley, 2003; Percy and Gründling, 2014; Rismondo et al., 2021a). The modification of TAs with d-alanine residues leads to a reduction in the negative surface charge and affects the activity of peptidoglycan hydrolases (Brown et al., 2013; Tesson et al., 2022; Vadyvaloo et al., 2004). Furthermore, lack of d-alanine modifications of TAs was shown to increase lysozyme sensitivity in *Staphylococcus aureus* (Herbert et al., 2007). Interestingly, the cell surface of the *eslB* mutant is more negatively charged, similar to that of a strain lacking d-alanine modifications on TAs. Based on our results, we propose the following model: The absence of EslB seems to affect the production and/or distribution of UDP-GlcNAc between different pathways, leading to the production of a thinner peptidoglycan layer. The reduced peptidoglycan thickness could subsequently result in the presentation of a larger portion of WTA on the bacterial cell surface, which would explain the higher negative surface charge (Fig. 9). This higher negative surface charge would enhance the binding capability and/or activity of cationic antimicrobial peptides, lysozyme as well as peptidoglycan hydrolases (Low et al., 2011; Neuhaus and Baddiley, 2003; Ragland and Criss, 2017; Steudle and Pleiss, 2011; Weidenmaier et al., 2003) and furthermore explain the sensitivity of the *eslB* mutant towards CwlO activity, lysozyme and cationic antimicrobial peptides (Burke et al., 2014; Rismondo et al., 2021b).

**Figure 9:**
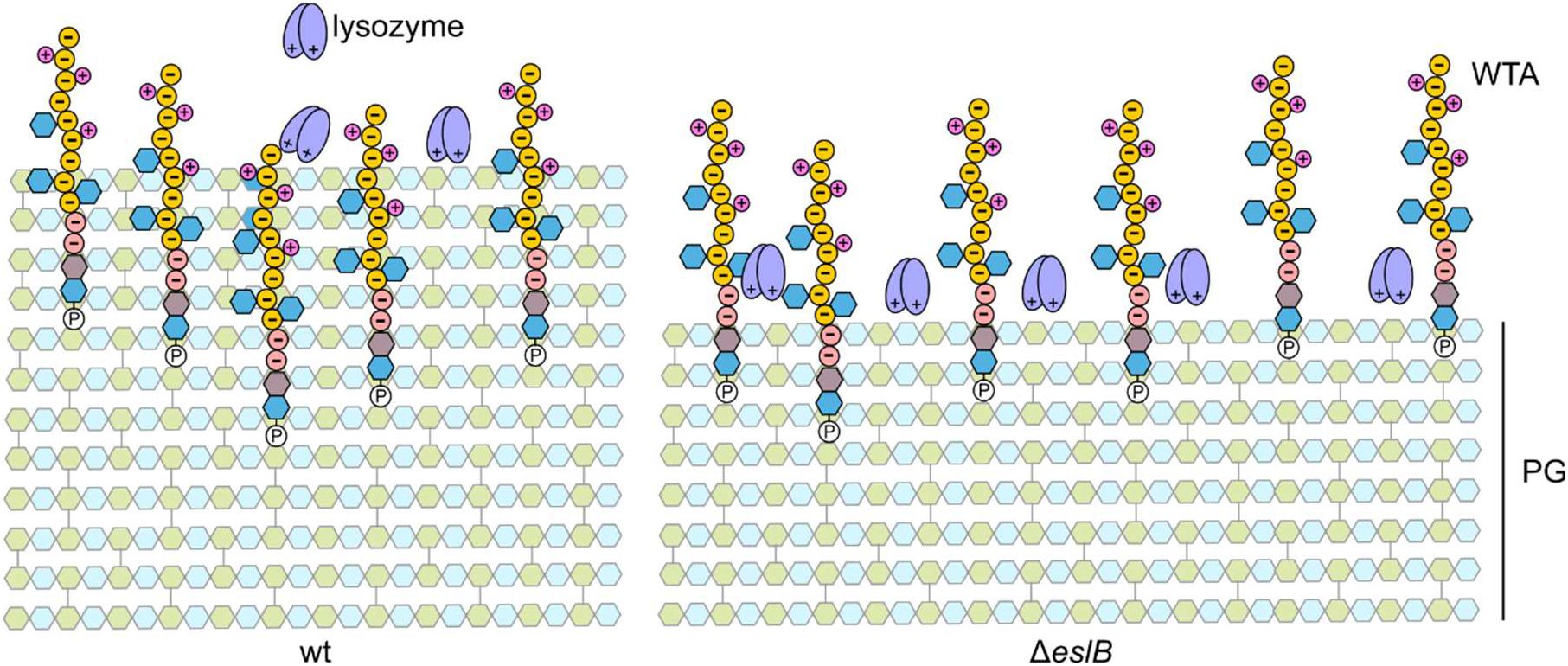
Model of altered WTA presentation on the cell surface of an *eslB* mutant. The Gram-positive cell wall is composed of a thick layer of peptidoglycan (PG) and wall teichoic acids (WTA), which are attached to the Mur*N*Ac-moiety of the PG backbone (Brown et al., 2013). WTA of *L. monocytogenes* 10403S are composed of a negatively charged ribitol phosphate backbone (indicated by -), which can be modified with GlcNAc residues (blue hexagons) and positively charged d-alanine residues (indicated by +). The absence of EslB seems to affect the production and/or distribution of the PG precursor UDP-Glc*N*Ac between different pathways, which results in the production of a thinner PG layer. WTAs are thought to partially stick out of the PG layer and we propose that due to the thinner PG layer produced by the *eslB* mutant, a larger portion of the WTA backbone, which is negatively charged, could be presented on the bacterial cell surface. This would result in a higher negative cell surface charge of the *eslB* mutant, which would affect the binding capability and/or activity of PG hydrolases, lysozyme and cationic antimicrobial peptides (Low et al., 2011; Ragland and Criss, 2017; Steudle and Pleiss, 2011; Weidenmaier et al., 2003).

The essential two-component system WalRK stimulates the transcription of several peptidoglycan hydrolases in Gram-positive bacteria (Delaune et al., 2011; Delauné et al., 2012; Dubrac et al., 2008; Dubrac and Msadek, 2004; Howell et al., 2003). In *B. subtilis*, WalRK also regulates the expression of genes involved in WTA biosynthesis and export (Howell et al., 2003). The regulon of the WalRK system has not yet been determined for *L. monocytogenes*, however, it was shown that the system is essential (Fischer et al., 2022). Inactivation of WalRK usually leads to cell death of wildtype cells due to loss of peptidoglycan hydrolase activity, however, cell death could be prevented by the inhibition of peptidoglycan biosynthesis (Salamaga et al., 2021). We have isolated several *eslB* suppressors with mutations in *walR* and *walK*. As it is unlikely that all of these mutations are gain-of-function mutations, we hypothesize that these mutations result in a reduced activity of the WalRK system. This would lead to a reduced peptidoglycan hydrolase activity as well as a reduction in WTA content, and reduction of the latter would at least partially restore the bacterial cell surface charge of the *eslB* mutant.

Taken together, we could show that the lack of EslB results in a defect in peptidoglycan biosynthesis, which can be suppressed by modulating the activity of enzymes involved in either peptidoglycan biosynthesis or hydrolysis. Our results suggest that the production or distribution of the peptidoglycan precursor UDP-GlcNAc between different pathways might be disturbed in the *eslB* mutant. Further studies are required to prove this hypothesis and to determine the function of the ABC transporter EslABC.

## AUTHOR CONTRIBUTION STATEMENT

**Lisa Maria Schulz:** Conceptualization, Funding acquisition, Investigation, Data analysis, Visualization, Writing – review & editing. **Patricia Rothe:** Investigation. **Sven Halbedel:** Supervision, Writing – review & editing. **Angelika Gründling:** Conceptualization, Funding acquisition, Writing – review & editing. **Jeanine Rismondo:** Conceptualization, Funding acquisition, Investigation, Data analysis, Visualization, Writing – original draft preparation.

## ACKNOWLEDGEMENTS

We thank Ivan Andrew and Jaspreet Haywood from the CSC Genomics Laboratory, Hammersmith Hospital, for their help with the whole genome sequencing and Annika Gillis for help with genome sequence analysis. We also thank Julia Busse for technical assistance and Tayfun Acar for the help with some experiments. We are grateful to Prof. Jörg Stülke for providing JR and LMS with laboratory space, equipment and consumables and to the Göttingen Center for Molecular Biosciences (GZMB) for financial support. This work was funded by the Wellcome Trust grant 210671/Z/18/Z and MRC grant MR/P011071/1 to AG, the German research foundation (DFG) grants RI 2920/1-1, RI 2920/2-1 and RI 2920/3-1 to JR, HA 6830/4 to SH. LMS was supported by the Göttingen Graduate School for Neurosciences, Biophysics, and Molecular Biosciences (GGNB, DFG grant GSC226/4).

