## Supplemental Material for "Imbalance of peptidoglycan biosynthesis alters the cell surface charge of *Listeria monocytogenes*"

### SUPPLEMENTAL TABLES

**Table S1: Bacterial strains used in this study**

| Unique ID | Strain name and resistance | Source |
|---|---|---|
| ***Escherichia coli* strains** | | |
| ANG1264 | DH5α pKSV7; AmpR | (Smith and Youngman, 1992) |
| ANG4242 | XL1-Blue pIMK2; KanR | (Monk et al., 2008) |
| ANG4243 | XL1-Blue pIMK3; KanR | (Monk et al., 2008) |
| ANG4236 | XL1-Blue pKSV7-Δ*eslB*; AmpR | (Rismondo et al., 2021) |
| EJR50 | XL1-Blue pAC7-$P_{dlt}$; AmpR | This study |
| EJR51 | XL1-Blue pAC7-$P_{dlt}$*; AmpR | This study |
| EJR52 | XL1-Blue pIMK3-*murA*; KanR | This study |
| EJR58 | S17-1 pIMK3; KanR | This study |
| EJR59 | S17-1 pIMK3-*murA*; KanR | This study |
| EJR63 | XL1-Blue pKSV7-Δ*cwlO*; AmpR | This study |
| EJR106 | XL1-Blue pIMK3-*glmR*; KanR | This study |
| EJR107 | XL1-Blue pIMK3-*glmU*; KanR | This study |
| EJR108 | XL1-Blue pIMK3-*glmM*; KanR | This study |
| EJR112 | XL1-Blue pIMK3-*glmS*; KanR | This study |
| EJR114 | XL1-Blue pIMK3-*cwlO*; KanR | This study |
| EJR115 | S17-1 pIMK3-*cwlO*; KanR | This study |
| EJR132 | S17-1 pIMK3-*glmR*; KanR | This study |
| EJR133 | S17-1 pIMK3-*glmU*; KanR | This study |
| EJR134 | S17-1 pIMK3-*glmM*; KanR | This study |
| EJR135 | S17-1 pIMK3-*glmS*; KanR | This study |
| ***Listeria monocytogenes* strains** | | |
| | EGD-e | (Glaser et al., 2001) |
| ANG1263 | 10403S; StrepR | (Bishop and Hinrichs, 1987) |
| ANG4275 | 10403SΔ$eslB_{(1)}$; StrepR | (Rismondo et al., 2021) |

| | | |
|---|---|---|
| ANG4688 | 10403SΔ*eslB*$_{(1)}$ pIMK3-*eslB* (or short: 10403SΔ*eslB*$_{(1)}$ compl.); StrepR KanR | (Rismondo et al., 2021) |
| ANG5386 | 10403SΔ*eslB*$_{(1)}$ *murZ*$^{M240fs}$; StrepR | This study |
| ANG5479 | 10403SΔ*eslB*$_{(1)}$ *walK*$^{R553H}$; StrepR | This study |
| ANG5480 | 10403SΔ*eslB*$_{(1)}$ *walK*$^{N283E}$; StrepR | This study |
| ANG5488 | 10403SΔ*eslB*$_{(1)}$ *ftsX*$^{G253R}$; StrepR | This study |
| ANG5489 | 10403SΔ*eslB*$_{(1)}$ *walK*$^{A570V}$; StrepR | This study |
| ANG5499 | 10403SΔ*eslB*$_{(1)}$ *reoM*$^{K23fs}$; StrepR | This study |
| ANG5662 | 10403SΔ*eslB*$_{(2)}$; StrepR | (Rismondo et al., 2021) |
| ANG5663 | 10403SΔ*eslB*$_{(2)}$ pPL3e-P$_{eslA}$-*eslABC* (or short: 10403SΔ*eslB*$_{(2)}$ compl.); ErmR StrepR | (Rismondo et al., 2021) |
| ANG5685 | 10403SΔ*eslB*$_{(3)}$; StrepR | This study |
| ANG5698 | 10403SΔ*eslB*$_{(1)}$ *murZ*$^{Q307fs}$; StrepR | This study |
| ANG5699 | 10403SΔ*eslB*$_{(2)}$ *walK*$^{I583T}$; StrepR | This study |
| ANG5708 | 10403SΔ*eslB*$_{(2)}$ *prpC*$^{P159L}$; StrepR | This study |
| ANG5710 | 10403SΔ*eslB*$_{(2)}$ *cwlO*$^{R106H}$; StrepR | This study |
| ANG5714 | 10403SΔ*eslB*$_{(2)}$ *ftsE*$^{Q220-}$; StrepR | This study |
| ANG5717 | 10403SΔ*eslB*$_{(2)}$ *pbpA1*$^{G125D}$; StrepR | This study |
| ANG5729 | 10403SΔ*eslB*$_{(3)}$ *tarL*$^{P282L}$; StrepR | This study |
| ANG5730 | 10403SΔ*eslB*$_{(3)}$ *cwlO*$^{R135P}$; StrepR | This study |
| ANG5733 | 10403SΔ*eslB*$_{(2)}$ *walR*$^{S216G}$; StrepR | This study |
| ANG5734 | 10403SΔ*eslB*$_{(2)}$ *walK*$^{H463D}$; StrepR | This study |
| ANG5737 | 10403SΔ*eslB*$_{(2)}$ *walK*$^{V234L}$; StrepR | This study |
| ANG5741 | 10403SΔ*eslB*$_{(2)}$ *walK*$^{R480H}$; StrepR | This study |
| ANG5746 | 10403SΔ*eslB*$_{(3)}$ $P_{dlt}$*; StrepR | This study |
| ANG5749 | 10403SΔ*eslB*$_{(3)}$ *walK*$^{H368Y}$; StrepR | This study |
| ANG5750 | 10403SΔ*eslB*$_{(3)}$ *walR*$^{E12G}$; StrepR | This study |
| ANG5754 | 10403SΔ*eslB*$_{(3)}$ *walK*$^{P600S}$; StrepR | This study |
| LJR24 | 10403S pIMK3; StrepR KanR | This study |
| LJR25 | 10403SΔ*eslB*$_{(2)}$ pIMK3; StrepR KanR | This study |
| LJR26 | 10403S pIMK3-*murA*; StrepR KanR | This study |
| LJR27 | 10403SΔ*eslB*$_{(2)}$ pIMK3-*murA*; StrepR KanR | This study |
| LJR37 | 10403SΔ*cwlO*; StrepR | This study |
| LJR56 | 10403SΔ*eslB*$_{(3)}$ pPL3e-P$_{eslA}$-*eslABC* (or short: 10403SΔ*eslB*$_{(3)}$ compl.); ErmR StrepR | This study |
| LJR63 | 10403SΔ*eslB*$_{(2)}$ pIMK3-*glmR*; StrepR KanR | This study |

| | | |
|---|---|---|
| LJR64 | 10403S$\Delta$*eslB*$_{(2)}$ pIMK3-*glmU*; StrepR KanR | This study |
| LJR65 | 10403S$\Delta$*eslB*$_{(2)}$ pIMK3-*glmM*; StrepR KanR | This study |
| LJR66 | 10403S$\Delta$*eslB*$_{(2)}$ pIMK3-*glmS*; StrepR KanR | This study |
| LJR103 | 10403S$\Delta$*cwlO* pIMK3-*cwlO*; StrepR KanR | This study |
| LJR114 | 10403S$\Delta$*cwlO* pIMK3-*cwlO* pKSV7-$\Delta$*eslB*; StrepR KanR Cam | This study |
| LJR119 | 10403S$\Delta$*cwlO* pIMK3-*cwlO* $\Delta$*eslB*; StrepR KanR | This study |
| LMJR138 | EGD-e $\Delta$*clpC* | (Rismondo et al., 2016) |
| ***Bacillus subtilis* strains** | | |
| | 168 *trpC2* | (Burkholder and Giles, 1947) |
| BLMS4 | 168 *trpC2 amyE*::P$_{dlt}$-*lacZ* | This study |
| BLMS5 | 168 *trpC2 amyE*::P$_{dlt}^{*}$-*lacZ* | This study |

**Table S2: Primers used in this study**

| Number | Name | Sequence |
|---|---|---|
| JR90 | p3-*murA* fw *Nco*I | CATGCCATGGAAAAAATTATTGTACGCGGTGGAAA |
| JR91 | p3-*murA* rev *Sal*I | ACGCGTCGACTTAGAATAAAGACGCTAAGTTTGTTAC |
| JR134 | p3-*glmR* fw *Nco*I | CATGCCATGGGGAAAAAGGAAACGAAACCTAGAGTAG |
| JR135 | p3-*glmR* rev *Sal*I | ACGCGTCGACTCACTCCTTTTCATTAGTTTGGTTA |
| JR136 | p3-*glmU* fw *Nco*I | CATGCCATGGGGTCAAAACGATATGCTGTAGTGC |
| JR137 | p3-*glmU* rev *Sal*I | ACGCGTCGACTTATTTACCATGATTCAAATGTTTCG |
| JR138 | p3-*glmM* fw *Nco*I | CATGCCATGGGTAAATATTTCGGTACGGATGG |
| JR139 | p3-*glmM* rev *Sal*I | ACGCGTCGACTTAATCGTTAAGTGCCATTTCTGAA |
| JR140 | p3-*glmS* fw *BamH*I | GCGGGATCCCTGTGGAATCGTTGGATATATAGG |
| JR141 | p3-*glmS* rev *Xma*I | TCCCCCCGGGTTATTCTACTGTGACACTTTTAGC |
| LMS93 | *dlt* promoter fw | CGCGGATCCTTCATAAATCCACCTCTTCATTGTTTCATTT |
| LMS94 | *dlt* promoter rev | CCGGAATTCGTCCAACCTCTTTTCTAGCATTAAAAC |
| LMS104 | *cwlO* down fw | ATTGCGATCTCACTTGTAGGATATGGCCGCGTAG |
| LMS105 | *cwlO* down rev | CGCGGATCCGTAGCTTCATTTGATGACGAACTTG |
| LMS106 | *cwlO* up fw | CGGGGTACCGAGCAGCAAGGAGAATATGGC |

| | | |
|---|---|---|
| LMS107 | *cwlO* up rev | ATATCCTACAAGTGAGATCGCAATAAACGTATTCT TTTTC |
| LMS226 | p3-*cwlO* fw *Nco*I | TTTCCATGGGGAAAAAGAATACGTTTATTGCGATC TCACT |
| LMS227 | p3-*cwlO* rev *BamH*I | AAAGGATCCTTAGAAGTTAGCTACGCGGCC |

### SUPPLEMENTAL FIGURES AND LEGENDS

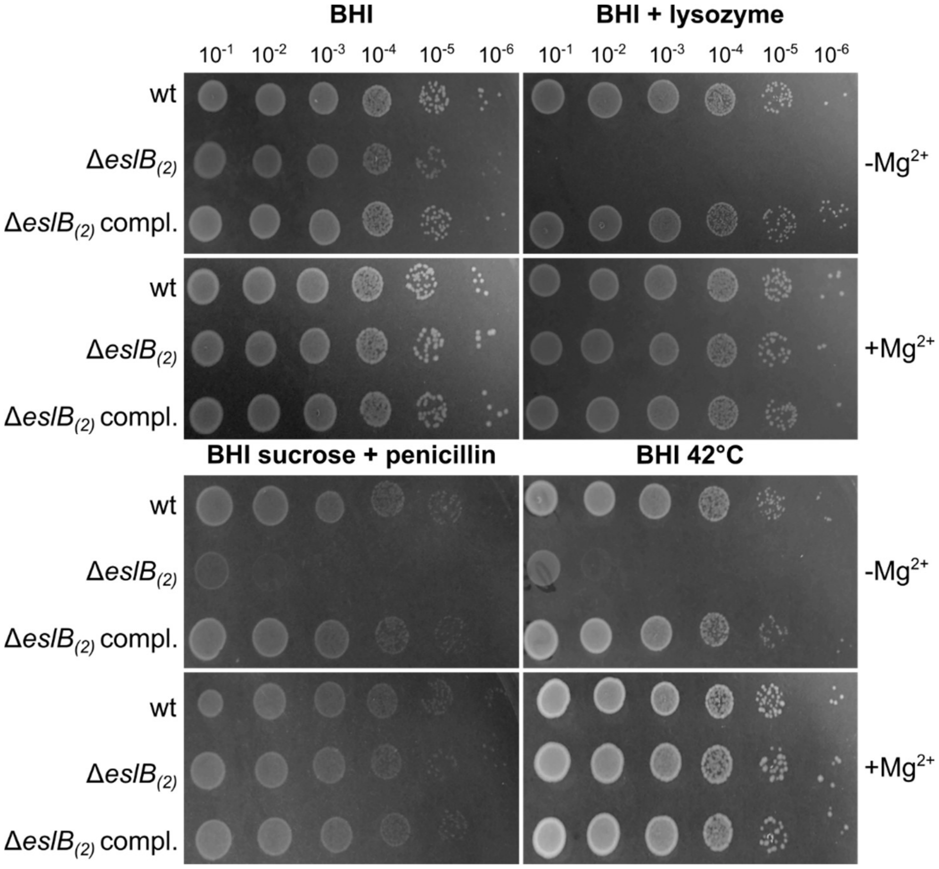


**Figure S1: Magnesium rescues growth deficits of the *eslB* mutant.** Drop dilution assay. Dilutions of *L. monocytogenes* strains 10403S (wt), Δ*eslB*$_{(2)}$ and Δ*eslB*$_{(2)}$ compl. were spotted on BHI plates, BHI plates containing 100 µg/ml lysozyme or containing 0.5 M sucrose and 0.025 µg/ml penicillin G and were incubated at 37°C, or on BHI plates that were incubated at 42°C. BHI plates contained 20 mM $Mg^{2+}$ were indicated.

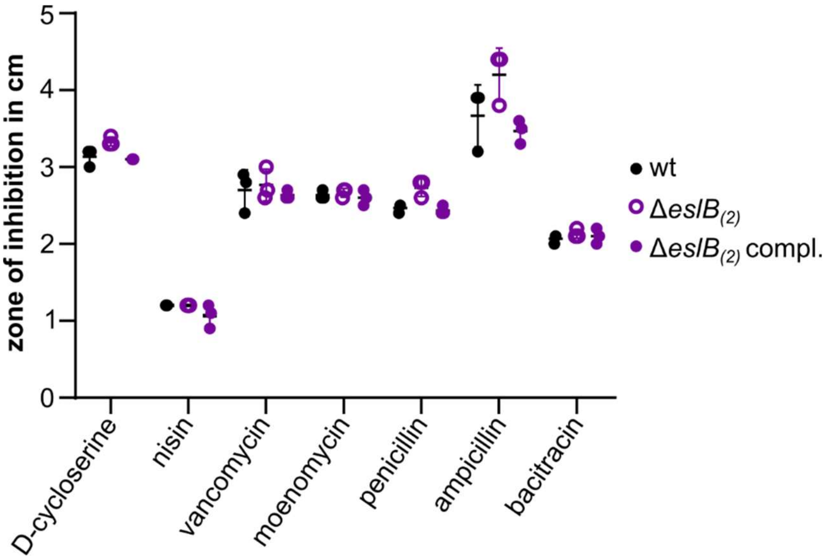


**Figure S2: Resistance of *L. monocytogenes* strains towards cell wall-targeting antibiotics.** Disk diffusion assay. *L. monocytogenes* strains 10403S (wt), Δ*eslB*$_{(2)}$ and Δ*eslB*$_{(2)}$ compl. were spotted on BHI plates. Antibiotic-soaked disks placed on top the agar surface and the plates incubated for 24h at 37°C. The inhibition zones for the indicated strains were measured and the average values and standard deviation of at least three independent experiments were plotted. For statistical analysis, a one-way ANOVA coupled with a Dunnett's multiple comparison test was used.

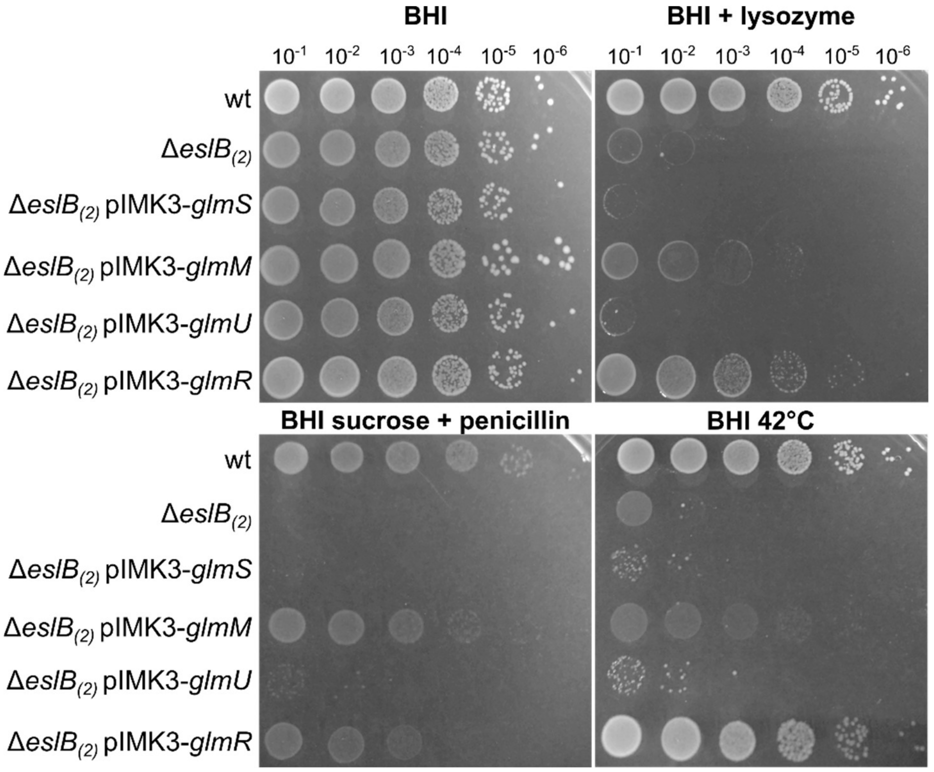


**Figure S3: Overexpression of *glmM* and *glmR* leads to partial suppression of *eslB* phenotypes.** Drop dilution assay. Dilutions of *L. monocytogenes* strains 10403S (wt), Δ*eslB*$_{(2)}$, Δ*eslB*$_{(2)}$ pIMK3-*glmS*, Δ*eslB*$_{(2)}$ pIMK3-*glmM*, Δ*eslB*$_{(2)}$ pIMK3-*glmU* and Δ*eslB*$_{(2)}$ pIMK3-*glmR*

were spotted on BHI plates, BHI plates containing 100 µg/ml lysozyme or containing 0.5 M sucrose and 0.025 µg/ml penicillin G and were incubated at 37°C or on BHI plates that were incubated at 42°C. Plates contained 1 mM IPTG to induce expression of *glmS*, *glmM*, *glmU* and *glmR*.

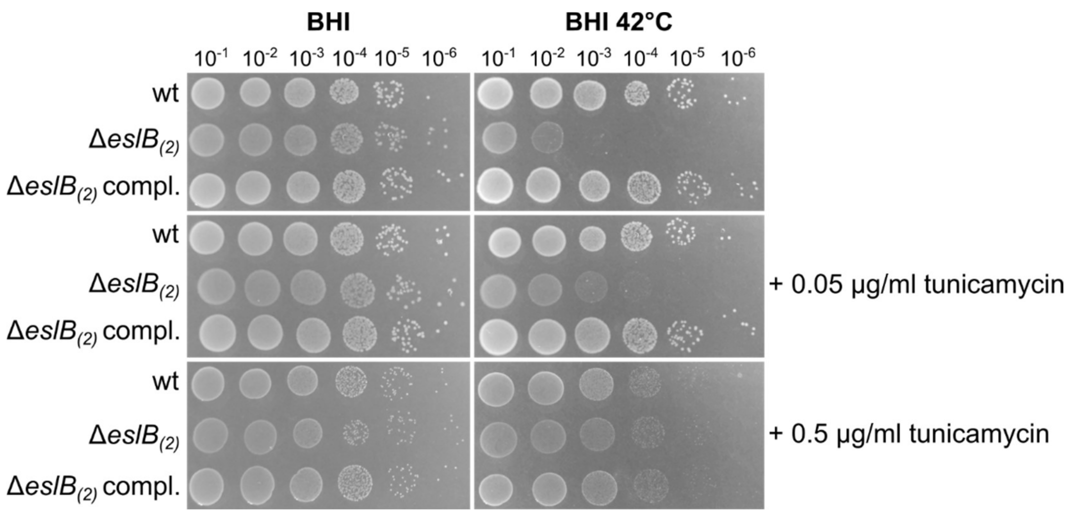


**Figure S4: Chemical inactivation of TarO partially rescues heat sensitivity of the *eslB* mutant.** Drop dilution assay. Dilutions of *L. monocytogenes* strains 10403S (wt), Δ*eslB*$_{(2)}$ and Δ*eslB*$_{(2)}$ compl. were spotted on BHI plates, BHI plates containing 0.05 or 0.5 µg/ml tunicamycin and were incubated at 37°C or 42°C.
